# Conserved neuropeptidergic regulation of intestinal integrity in invertebrate models of aging

**DOI:** 10.1101/2022.02.24.481867

**Authors:** Anupama Singh, Bhagyashree Kaduskar, Kirthi C Reddy, Caroline Kumsta, Ethan Bier, Malene Hansen, Sreekanth H Chalasani

## Abstract

Age-related decline in intestinal barrier function impacts survival across species, but the underlying cell intrinsic and extrinsic factors remain unclear. Here, we demonstrate a role for neuropeptides in regulating aging-associated increases in intestinal leakiness. Adult-specific knockdown of insulin-like peptides *daf-28* or *ins-7* in *C. elegans* neurons or *dilp3* in *D. melanogaster* neurons improves intestinal integrity and lifespan of these animals, respectively. Neuropeptide knockdowns activate intestinal DAF-16/FOXO transcription factor that likely increases expression of epithelial barrier components. Furthermore, age-associated changes in neuronal DAF-28 peptide secretion mirrors *C. elegans* reproductive span and overexpression of this peptide suppresses the improved intestinal health in long-lived germline-less animals. Collectively, we show that intestinal integrity is subject to neuronal regulation, and this neuropeptidergic axis may be modulated by the animal’s germline.

## Main

Animal aging involves a coordinated decline in function of various tissues. While individual tissues deteriorate at different rates^1^, communication between them can profoundly affect aging across species (reviewed in^2^). Consistently, an injured or diseased brain can trigger intestinal leakiness and reduce gastric motility, two characteristic features of intestinal aging, in both vertebrate and invertebrates^3-5^. Evidence of the role of inter-tissue signaling in animal aging is also seen in studies where transferring circulating polypeptides GDF11, myostatin and osteocalcin from young to older rodents has a rejuvenating effect on the latter^6-9^. However, little has been done to study whether age-associated changes in one tissue actively modulates aging in other distal tissues – an essential link in aging across species.

In *C. elegans*, multiple conserved lifespan-modifying signaling pathways have been shown to function primarily in neurons, or their intestines, among other tissues^10-14^. Also, altering synaptic transmission from motor neurons or specific chemosensory neurons can improve animal lifespan^15-19^, confirming a role for neuronal signaling in animal aging. Similarly, their intestine has also been shown to play a crucial role in animal health and survival^11,20-22^. Despite this less is known about how changes in neuronal signaling affects aging in distal tissues like the intestine. Of the many age-associated changes in the intestine, barrier dysfunction is conserved between humans and laboratory animals^23,24^. This barrier dysfunction phenotype is readily quantified as the extent of intestinal leakiness in so-called “Smurf” assays. Here, the localization of a blue-dye co-ingested with food is monitored; the animal turns blue when its intestinal barrier is compromised, resulting in the dye spreading from its intestine into the body cavity^21,23,25^. Along with *C. elegans*, we also used the fruit fly *D. melanogaster* to probe the role of inter-tissue signals in affecting intestinal barriers in aged animals.

Here, we leveraged neuron-selective RNA interference (RNAi) method to alter neuronal signaling and measured its impact on the aging outcomes in the intestine. Using *C. elegans* and *D. melanogaster* in this neuron-focused approach, we show a conserved role for neuronal peptides in modifying intestinal integrity during aging. We identify the specific neuronal peptides and their intestinal receptors that regulate age-related changes in intestinal leakiness in both nematodes and flies. Next, we show that blocking neuropeptide signaling, non-cell autonomously activated DAF-16/FOXO in the intestine. Additionally, DAF-16/FOXO has been shown to directly bind promoters of several epithelial barrier components, linking neuropeptide signaling with intestinal barrier function. Finally, we gained insights into the timing of this communication, and link reproductive activity with neuron-to-intestine insulin-like peptide signals and intestinal aging. Moreover, both *C. elegans* and *Drosophila* intestine bears many similarities to the human gastrointestinal tract^26,27^, implying that our findings could be conserved across species.

## Results

### Neuropeptide signaling selectively affects age-associated decline in intestinal integrity

To test whether *C. elegans* neurons can influence age-associated changes in other tissues, we knocked down *unc-13/Munc-13* (synaptic vesicle-priming for neurotransmitter release^28^), *snb-1/synaptobrevin* (synaptic-vesicle associated vSNARE^29^) or *unc-31/CAPS* (activator protein for peptide secretion^30^) selectively in neurons of young adults (i.e., day 1 as reproductive adults) and monitored age-associated changes in other tissues (**Fig. 1A**). We first assessed intestinal health by measuring age-related increase in intestinal leakiness using the Smurf assay. Briefly, aged animals (day 15 of adulthood) were fed a blue dye mixed with bacteria for 3 hours, after which the dye-spread inside the body was monitored. We further evaluated intestinal leakiness by combining fluorescein with the blue dye, allowing us to obtain a more quantitative estimate of intestinal leakiness, besides the binary classification into Smurfs and non-Smurfs (see Methods). Neuron-specific feeding-RNAi was implemented using transgenic animals expressing *sid-1* cDNA (SID-1 is membrane-associated, double-stranded RNA transporter), in a *sid-1(qt9)* null mutant^31^, using a pan-neuronal *rgef-1* promoter (Y. Yang, C. Kumsta and M. Hansen, manuscript in preparation). We observed that neuronal *unc-31/CAPS* (**Fig. 1B, Supplementary Table 1**), but not *unc-13/Munc-13* or *snb-1/synaptobrevin* (**Extended DataFig. 1A, Supplementary Table 1**) knockdowns reduced the age-related increase in intestinal leakiness, a phenotype we refer to as “HypoSmurfs”. Next, we tested whether neuropeptides also affected other intestinal functions like defecation and gastric motility. We found that neuronal *unc-31/CAPS* knockdowns had no effect on age-associated declines in defecation (**Extended Data Fig. 1B**) or gastric motility (**Extended Data Fig. 1C**). In addition, knocking down *unc-13/Munc-13, snb-1/synaptobrevin*, or *unc-31/CAPS* in neurons had no significant effect on the age-dependent declines in the rates of both body bends and pharyngeal pumping (**Extended Data Fig. 1D-E**). These results highlight the specificity of neuropeptides in selectively regulating intestinal barrier integrity (Smurf phenotype), without impacting age-associated changes in defecation, gastric motility, pharyngeal pumping, or body movements.

**Fig. 1.**
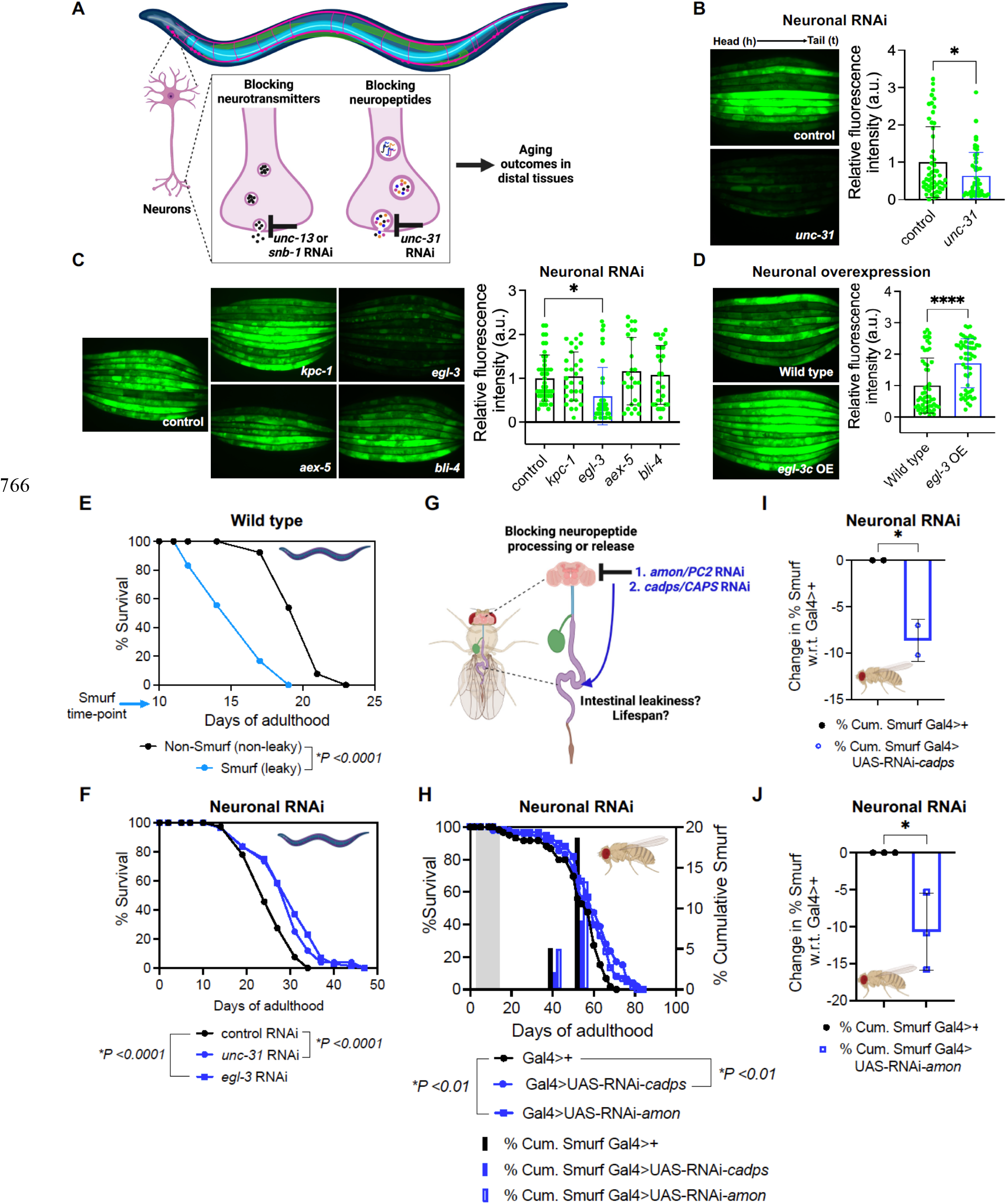
Blocking neuropeptides delays age-related intestinal leakiness in adult *C. elegans* and *D. melanogaster* models. **(A)** Schematic of experimental approach of reducing neurotransmitter and neuropeptide release in neurons to assess their effects on aging phenotypes in multiple *C. elegans* tissues. **(B)** FluoroSmurf phenotype of day 15 old animals during neuron-specific knockdown *unc-31/CAPS* feeding-RNAi, quantified n=2, N=54 (control), and N=48 (*unc-31/CAPS)*, out of 8 trails, * *P<0*.*05*, two-tailed t-test. Nematodes are aligned head (left) to tail (right), if not otherwise mentioned. Each dot represents relative fluorescent intensity in arbitrary units (a.u.) quantified from entire body of each animal, images acquired at constant exposure time. **(C)** FluoroSmurf phenotype of the four *C. elegans* pro-protein convertases, during their neuron-specific knockdown. Represented data quantified from n=1, N=45 (control), N=29 (*kpc-1/PC1)*, N=36 (*egl-3/PC2)*, N=26 (*aex-5/PC3)* and N=28 (*bli-4/PC4)*, out of 7 trails. * *P<0*.*05, one-way ANOVA* with Tukey’s multiple comparisons test. **(D)** The FluoroSmurf phenotype of transgenic animals pan-neuronally overexpressing *egl-3/PC2* (OE). Represented data quantified from n=1, N=56 (wild type), N=58 (*egl-3* OE), out of 5 trails. **** *P<0*.*0001*, two-tailed t-test. For (B-D), also see **Supplementary Table 1** for data of independent trials. **(E)** Lifespan of non-Smurfs (non-leaky animals) compared to Smurfs (leaky intestine), classified after day10 Smurf assay in Wild-type (N2-CGC) animals (* *P<0*.*0001*, log-rank test). Data is representative of two independent trials. See **Supplementary Table 2** for lifespan statistics. **(F)** Lifespans of neuron-specific knockdown of *unc-31/CAPS* or *egl-3/PC2* from day 1 of adulthood (* *P<0*.*0001*, log-rank test). Data is representative of more than three independent trials. See **Supplementary Table 1** for lifespan statistics. **(G)** Schematic of experimental approach for neuron-specific RNAi of *cadps/CAPS* or *amon/PC2*, to assess lifespan and intestinal leakiness in the *Drosophila*. **(H)** Smurf phenotype and lifespans of neuron-specific RNAi of *cadps/CAPS* or *amon/PC2* in flies (* *P<0*.*01*, log-rank test for lifespan). Grey bar represents duration (∼10 days) of neuron-specific GAL4 activation at 29ºC. **(I)** Reduced % Smurf in old flies (∼55 days), during neuronal knockdown of *cadps/CAPS* compared to Gal4>+ control. Error bars are std. dev. of n=2. * *P<0*.*05*, two-tailed t-test. **(J)** Reduced % Smurf in old flies (∼55 days), during neuronal knockdown of *amon/PC2* compared to Gal4>+ control. Error bars are std. dev. of n=3. * *P<0*.*05*, two-tailed t-test. See **Supplementary Table 3** for additional data of independent trials. *Also see Extended Data Fig. 1-2*.

To further investigate the role of neuropeptides in affecting intestinal integrity, we tested neuronal knockdowns of various components of the upstream peptide processing pathway, which is conserved from *C. elegans* to humans. Briefly, a pro-peptide is first synthesized and cleaved by one of four pro-protein convertases (i.e., KPC-1/PC1, EGL-3/PC2, AEX-5/PC3, and BLI-4/PC4) to generate peptide fragments. Basic amino acids from these peptide fragments are further removed by EGL-21/Carboxypeptidase E. The resulting peptides are then amidated by monooxygenase PAMN-1 and amidating lyase PGAL-1, before being packaged into dense core vesicles (DCVs) and released in an UNC-31/CAPS-dependent manner (**Extended Data Fig. 2A**)^32^. We found that knocking down the pro-protein convertase *egl-3/PC2*, but not *kpc-1/PC1, aex-5/PC3*, or *bli-4/PC4* in neurons attenuated aging-induced increases in intestinal leakiness (**Fig. 1C**). In addition, neuronal knockdown of *sbt-1/7B2*, the *egl-3/PC2* chaperone^33^ (**Extended Data Fig. 2B**) also reduced aging-induced intestinal leakiness, confirming a role for EGL-3/PC2 processing in regulating intestinal integrity. In addition, we also found that neuronal knockdown of the downstream *egl-21*/*Carboxypeptidase E* (**Extended Data Fig. 2C**) similarly affected intestinal aging, suggesting an intestinal integrity-modifying role for EGL-21 peptidase. Consistent with these findings, over-expression of *egl-3/PC2* in neurons increased intestinal leakiness in aged adults compared to controls, a phenotype we refer to as “HyperSmurfs” (**Fig. 1D**). However, knocking down the monooxygenase *pamn-1* or amidating lyase *pgal-1* in neurons had no effect on intestinal integrity in aged adults (**Extended Data Fig. 2D**), likely because of redundancy in the amidation step^34^. Collectively, these data suggest that *C. elegans* neurons use UNC-31/CAPS protein to release neuropeptides sequentially processed by EGL-3/PC2 and EGL-21/Carboxypeptidase E, whose function impairs intestinal integrity in aged animals.

We next tested whether differences in intestinal integrity were correlated with *C. elegans* lifespan, similar to what has been observed in flies^23^. On day 10 of adulthood, we separated animals based on their Smurf phenotype into leaky or non-leaky groups and monitored their lifespans independently. Animals with intact intestines (non-Smurfs) were longer lived (* *P<0*.*0001*, log-rank test) compared to animals with a leaky intestine (Smurfs), showing that intestinal integrity inversely correlates with lifespan (**Fig. 1E**)^25^. Consistently, we found that knocking down *unc-31/CAPS* and *egl-3/PC2* in neurons extended lifespan by ∼15% (* *P<0*.*0001*, log-rank test) and ∼20% (* *P<0*.*0001*, log-rank test), respectively (**Fig. 1F, Supplementary Table 2**), confirming that blocking neuropeptide processing or release improves intestinal integrity and lifespan.

Next, we examined whether effects of neuropeptide signaling on intestinal integrity and lifespan were conserved between nematodes and flies (**Fig. 1G**). We monitored intestinal leakiness and animal lifespan in neuron-specific RNAi knockdowns of the *Drosophila* CAPS homolog, *cadps*^35^ and PC2 homolog, *amon*^36^. We observed an ∼10% extension in lifespan (* *P<0*.*01*, log-rank test) (**Fig. 1H, Supplementary Table 3**) and reduced intestinal leakiness (**Fig. 1H-J, Supplementary Table 3**) in aged flies in both neuronal knockdowns, suggesting that Amon/PC2-processed neuropeptides, secreted in Cadps-dependent manner, negatively affect intestinal barrier function and lifespan. Collectively, these data show a conserved role for *C. elegans* and *D. melanogaster* homologs of CAPS and PC2 proteins in intestinal and animal aging.

### Insulin-like neuropeptides exacerbate age-related decline in intestinal integrity

While the *C. elegans* genome includes ∼150 neuropeptide genes that encode over 250 neuropeptides^32,37^, we focused our analysis on the 40 insulin-like peptides (ILPs) because of their known effects on growth, reproduction, and lifespan^38-41^. To identify candidate intestine-modifying ILPs, we knocked down individual insulin-like peptide genes in day 1 young adult neurons and monitored intestinal aging using Smurf assays (**Fig. 2A, S3A-D, Supplementary Table 4**). We found that neuronal knockdowns of two beta-type insulin-like peptides, *daf-28* and *ins-7*, consistently exhibited the strongest HypoSmurf phenotypes (**Fig. 2B-C, S3B, Supplementary Table 1**), identifying two intestine integrity-modifying neuropeptides. Consistent with this, neuronal over-expression of *daf-28* or *ins-7* reproducibly increased intestinal leakiness in aged adults (**Fig. 2D-E**), mimicking the HyperSmurf phenotype of neuronal *egl-3/PC2* over-expression (**Fig. 1D**). Further, neuron-specific knockdown of *daf-28 or ins-7* extended lifespan by *∼*15% (* *P<0*.*005*, log-rank test) and ∼30% (* *P<0*.*0001*, log-rank test) respectively (**Fig. 2F-G, Supplementary Table 2**). Together, our neuron-specific RNAi screen identified DAF-28 and INS-7 as the first insulin-like neuropeptides that function to antagonize intestinal integrity.

**Fig. 2.**
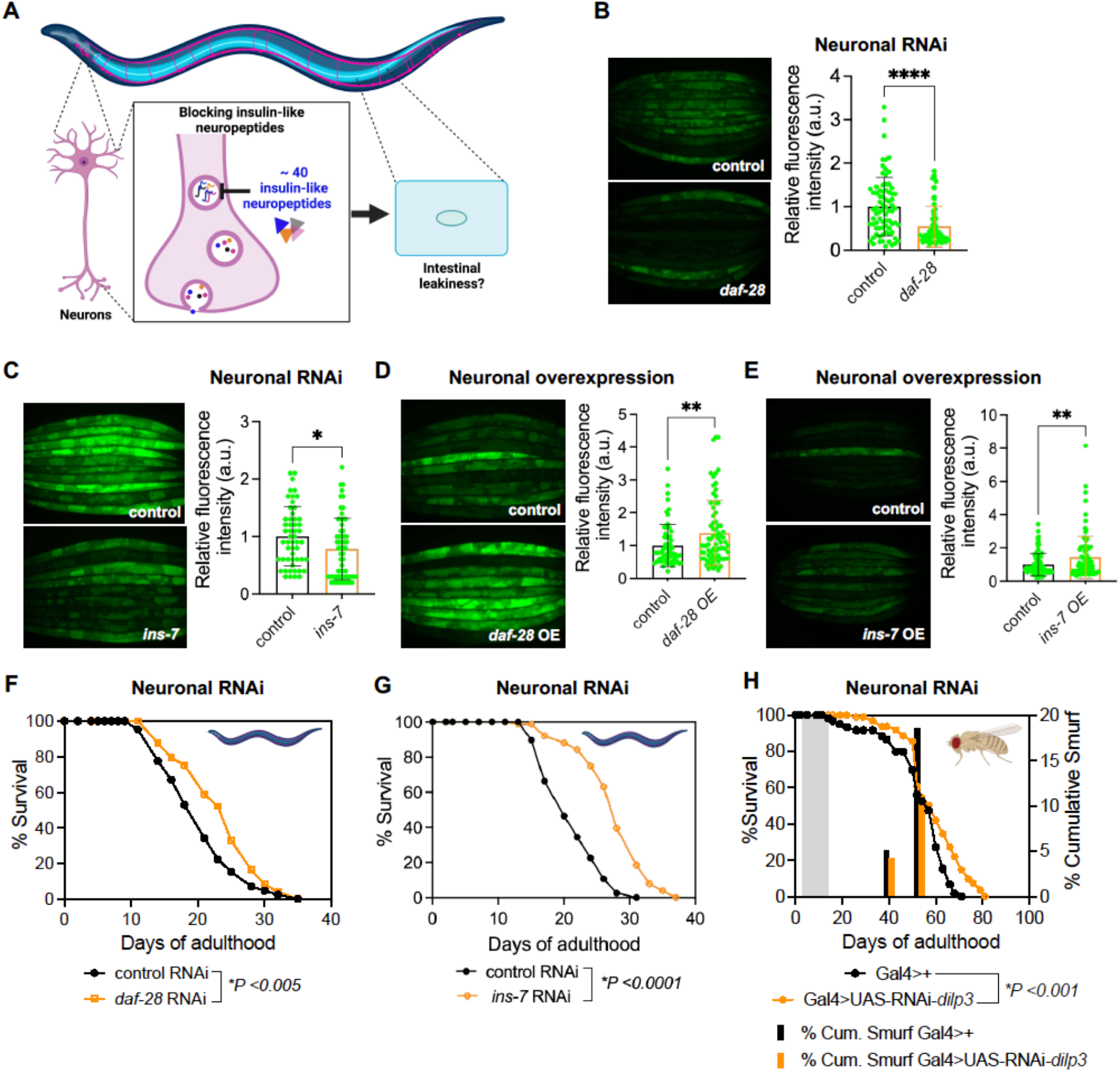
Conserved intestine-antagonizing role of insulin-like neuropeptides. **(A)** Schematic of experimental approach for neuron-specific knockdown of 35 (out of 40) insulin-like peptides to test their effect on intestinal leakiness in 15-day old *C. elegans*. **(B)** HypoSmurf phenotype of day 15 old animals during neuron-specific knockdown *daf-28*. Represented data quantified from n=1, N=76 (control), N=61 *(daf-28)*, out of 3 trails. **** *P<0*.*0001*, two-tailed t-test. **(C)** HypoSmurf phenotype of day 15 old animals during neuron-specific knockdown *ins-7*. Represented data quantified from n=1, N=53 (control), N=60 *(ins-7)*, out of 9 trials. * *P<0*.*05*, two-tailed t-test. **(D)** HyperSmurf phenotype of *C. elegans* pan-neuronally overexpressing *daf-28*. Represented data quantified from n=2, N=62 (control), N=75 (*daf-28* OE), out of 6 trails. ** *P<0*.*01*, two-tailed t-test. **(E)** HyperSmurf phenotype of *C. elegans* pan-neuronally overexpressing *ins-7*. Represented data quantified from n=2, N=89 (control), N=95 (*ins-7* OE), out of 4 trails. ** *P<0*.*01*, two-tailed t-test. For (B-E), also see **Supplementary Table 1** for data of independent trials. **(F)** Lifespan extension during neuron-specific knockdown of *daf-28 (*\* *P<0*.*005*, log-rank test), initiated from day 1 of adulthood. Data is representative of two independent trials. See **Supplementary Table 2** for lifespan statistics. **(G)** Lifespan extension during neuron-specific knockdown of *ins-7* (* *P<0*.*0001*, log-rank test), initiated from day 1 of adulthood. Data is representative of three independent trials. See **Supplementary Table 2** for lifespan statistics. **(H)** Neuron-specific RNAi of *dilp3* in flies reduced age-related intestinal leakiness and extended lifespan (* *P<0*.*001*, log-rank test for lifespan). Grey bar represents duration (∼10 days) of neuron specific GAL4 activation at 29ºC. See **Supplementary Table 3** for additional data of independent trials. *Also see Extended Data Fig. 3*.

We next tested whether *Drosophila* insulin-like peptides also play a similar intestine-modifying function. The fly genome encodes eight insulin-like peptides (Dilp1-8), and three of these (Dilp2, 3, and 5) are secreted from insulin producing cells (IPCs) in the brain {reviewed in^42,43^}. We found that knocking down *dilp3* (or *dilp2, dilp5*, see **Supplementary Table 3)** specifically in adult neurons reduced the age-related Smurf phenotype and improved lifespan by ∼12% (* *P<0*.*001*, log-rank test) (**Fig. 2H, Supplementary Table 3**). Taken together, these data suggest a conserved intestine-modifying role of insulin-like neuropeptides, affecting intestinal integrity and lifespan in both nematodes and flies.

### Neuropeptide signals affect cellular integrity via intestinal insulin signaling

In *C. elegans*, a single tyrosine kinase insulin/IGF-1-like receptor, DAF-2/InR, is known to integrate signals from all 40 insulin-like peptides (**Fig. 3A**)^44^. Moreover, both intestine-modifying neuropeptides, DAF-28 and INS-7, are known agonists of the DAF-2/InR^32,40^. We found that intestine-specific knockdown of *daf-2/InR* in day 1 young adults reduced intestinal leakiness on day 15 of adulthood (**Fig. 3B, Supplementary Table 1**), similar to what was observed when genes encoding insulin-like neuropeptides, enzymes for their processing, or components of the release machinery were reduced in neurons. Additionally, we found that intestine-specific knockdown of the downstream phosphoinositide 3-kinase, *age-1/PI3K*^45^ also reduced intestinal leakiness (**Extended Data Fig. 4A, Supplementary Table 1)**, indicating a cell-autonomous role for insulin signaling in impairing intestinal leakiness. Moreover, intestine-specific knockdown of *daf-2/InR* or *age-1/PI3K* extended lifespan by ∼60% (* *P<0*.*0001*, log-rank test) (**Fig. 3C, Supplementary Table 2)** and ∼40% (* *P<0*.*0001*, log-rank test) (**Extended Data Fig. 4B, Supplementary Table 2**), respectively, confirming inverse correlation between intestinal leakiness and animal lifespan. We next tested whether the single insulin receptor in *Drosophila*, dInR, had a similar effect on age-related changes in intestinal leakiness and lifespan. We observed that midgut-specific knockdown of *dInR* resulted in reduced intestinal leakiness and increased lifespan by ∼10% (* *P<0*.*001*, log-rank test), compared to controls (**Fig. 3D, Supplementary Table 3)**. Cumulatively, our data underscores a conserved cell-autonomous role of DAF-2/dInR signaling in impairing intestinal integrity and animal lifespan.

**Fig. 3.**
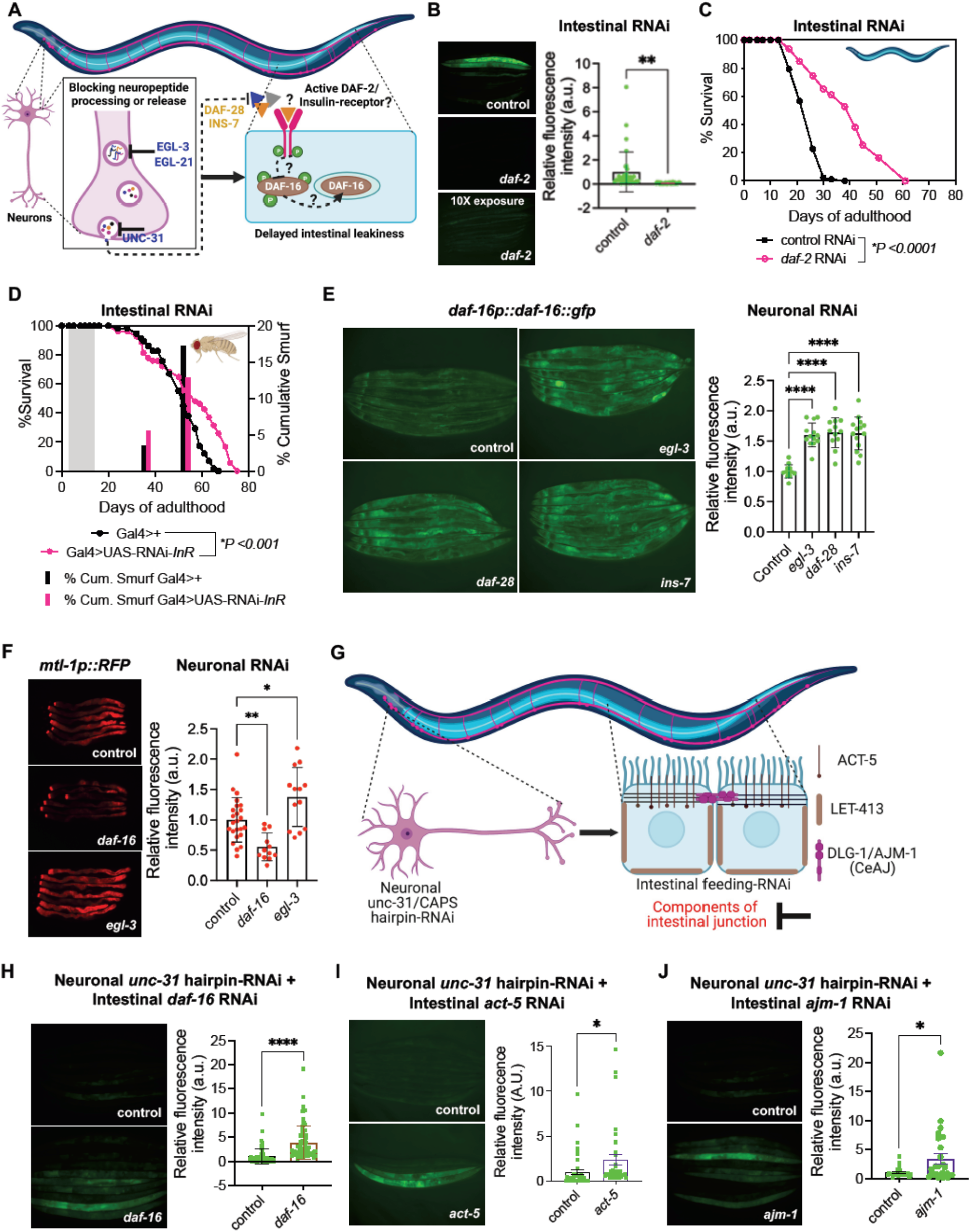
Reduced neuropeptide signals are integrated by intestinal insulin signaling and downstream epithelial barrier components. **(A)** Schematic depicting the working hypothesis to test role of intestinal insulin signaling and the downstream DAF-16/FOXO transcription factor, in converging neuropeptide signals to impact intestinal barrier. **(B)** HypoSmurf phenotype of intestine-specific KD of *daf-2/InR*. Represented data quantified from n=1, N=31 (control), N=29 (*daf-2/InR)*, out of 9 trails. ** *P<0*.*01*, two-tailed t-test. Lower panel image was acquired at 10X exposure time. See **Supplementary Table 1** for data of independent trials. **(C)** Lifespan extension during intestine-specific knockdown of *daf-2/InR (*\**P < 0*.*0001)*. RNAi was initiated on day 1 of adulthood. Data is representative of six independent trials. See **Supplementary Table 2** for lifespan statistics. **(D)** Midgut-specific RNAi of *dInR* reduced age-related intestinal leakiness and extended lifespan (\**P<0*.*001*) in flies. Grey bar represents duration (∼10 days) of midgut GAL4 activation at 29ºC. See **Supplementary Table 3** for additional data of independent trials. **(E)** DAF-16/FOXO intestinal expression monitored using *daf-16p::daf-16::gfp*, during neuron-specific RNAi of indicated genes initiated on day 1 and imaged on day 5. Error bars are std. dev. of N>12, **** *P<0*.*0001, one-way ANOVA* with Dunnett’s multiple comparisons test. Data is representative of three independent trials. **(F)** MTL-1 expression during neuron-specific RNAi of *egl-3/PC2 or daf-16/FOXO*. Error bars are std. dev. of N>12, ** *P<0*.*01*, * *P<0*.*05, one-way ANOVA* with Dunnett’s multiple comparisons test. Data is representative of two independent trials. **(G)** Schematic of experimental approach for testing the requirement of epithelial barrier components in regulating intestinal barrier integrity downstream of neuronal peptides. **(H)** Intestine-specific knockdown of *daf-16/FOXO* suppressed the HypoSmurf phenotype of neuronal *unc-31/CAPS* hairpin-RNAi transgenics. Represented data quantified from n=2, N=67 (control), N=75 *(daf-16)*, out of 5 trails. **** *P<0*.*0001*, two-tailed t-test. **(I)** Intestine-specific knockdown of *act-5* (actin) suppressed the HypoSmurf phenotype of neuronal *unc-31/CAPS* hairpin-RNAi transgenics. Represented data quantified from n=1, N=43 (control), N=36 *(act-5)*, out of 2 trails. * *P<0*.*05*, two-tailed t-test. **(J)** Intestine-specific knockdown of *ajm-1* (septate junction component) suppressed the HypoSmurf phenotype of neuronal *unc-31/CAPS* hairpin-RNAi transgenics. Represented data quantified from for n=1, N=24 (control), N=26 (*ajm-1)*. * *P<0*.*05*, two-tailed t-test. For (G-I), also see **Supplementary Table 1** for data of independent trials. *Also see Extended Data Fig. 4-5*.

**Fig. 4.**
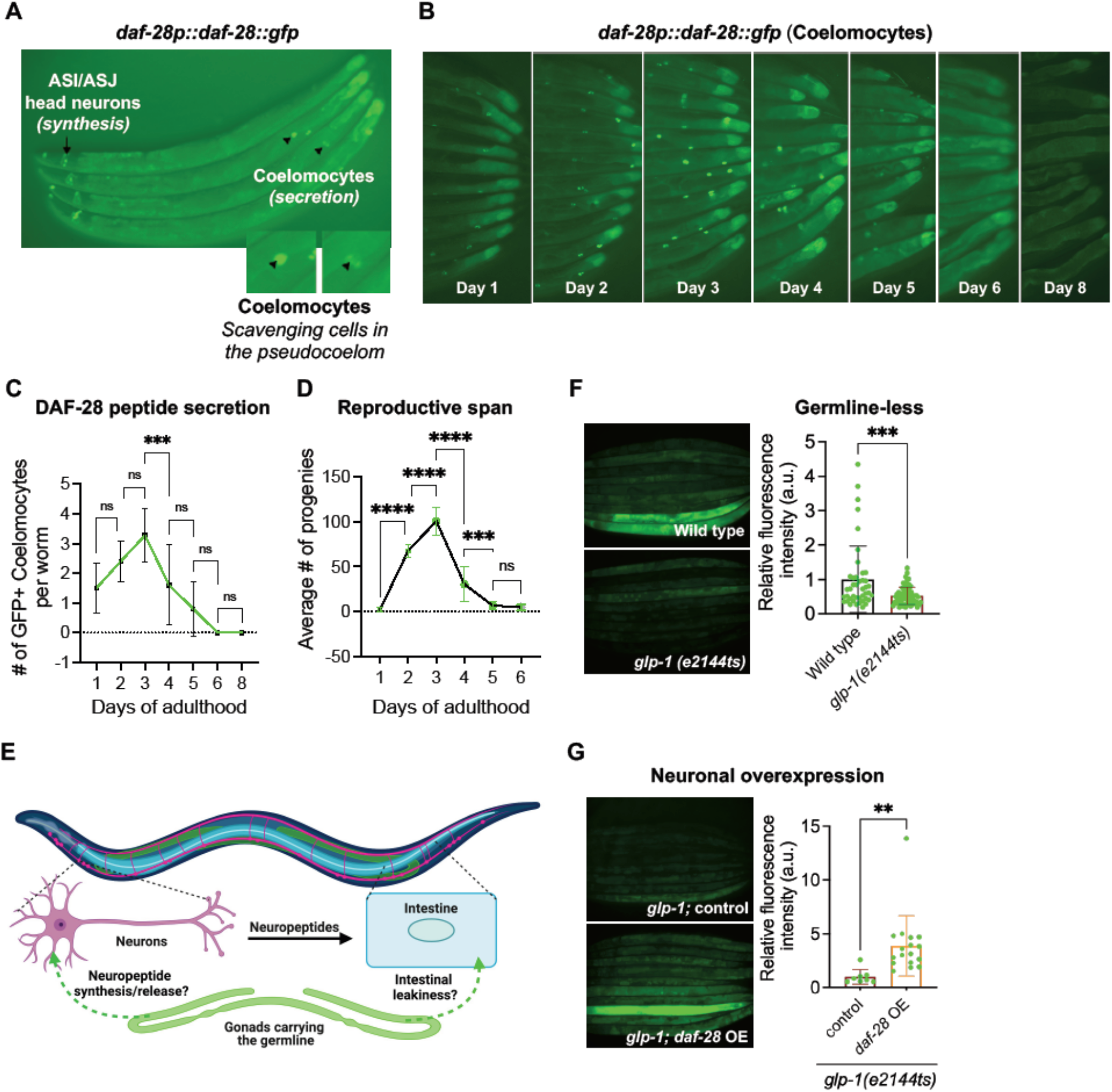
The age-related secretion of the intestine-modifying DAF-28 neuropeptide patterns reproductive span in *C. elegans*. **(A)** *daf-28p::daf-28::gfp* reporter used as a tool to monitor the age-related synthesis and secretion of the DAF-28 peptide. **(B-C)** The age-related secretion of DAF-28 neuropeptide measured as number of GFP-positive coelomocytes (scavenging cells). Error bars are std. dev. for N>10 \**\*\* P<0*.*001*, ns *P>0*.*05, one-way ANOVA* with Tukey’s multiple comparisons test. Data is representative of three independent trials. **(D)** Reproductive span of *daf-28p::daf-28::gfp* animals represented as average number of progenies produced each day of self-fertile period. Error bars are std. dev. for N=10, \*\**\*\* P<0*.*0001*, \**\*\* P<0*.*001*, ns *P>0*.*05, one-way ANOVA* with Tukey’s multiple comparisons test. Data is representative of two independent trials. **(E)** Schematic depicting experimental approach for testing role of germline in modulating the neuron-to-intestine crosstalks. **(F)** HypoSmurf phenotype of germline-less *glp-1(e2144ts)* mutants. Represented data quantified from n=1, N=43 (control), N=67 *(glp-1(e2144ts))*, out of 5 trails. *** *P<0*.*001*, two-tailed t-test. **(G)** Pan-neuronal overexpression of *daf-28* in *glp-1(e2144ts)* mutants suppressed their HypoSmurf phenotype. Represented data quantified from n=1, N=8 (*glp-1;* control), N=17 (*glp-1; daf-28* OE), out of 3 trails. ** *P<0*.*01*, two-tailed t-test. For (F-G), also see **Supplementary Table 1** for data of independent trials. *Also see Extended Data Fig. 6*.

Reduction in insulin signaling activates DAF-16/FOXO, localizing it to the nucleus^46^, enhancing DAF-16/FOXO promoter occupancy^47,48^ and transcriptional activation of a large number of genes^49,50^. We hypothesized that our neuron-specific knockdowns of insulin neuropeptides, or their processing and release machinery would result in reduced insulin signals and increased DAF-16/FOXO activation (**Fig. 3A**). To test this directly, we first monitored levels of DAF-16/FOXO using a *daf-16p:daf-16::gfp* translational reporter^51^. We found that neuron-specific knockdown of *egl-3/PC2, daf-28* or *ins-7* caused a significant increase in intestinal DAF-16/FOXO levels (**Fig. 3E**). Next, to validate the activation of DAF-16/FOXO, we monitored MTL-1 expression, a DAF-16/FOXO-regulated metallothionein protein, using an *mtl-1p::mtl-1::rfp* translational reporter^50^. We found that intestine-specific *daf-2/InR* knockdown upregulated MTL-1, while *daf-16/FOXO* knockdown suppressed its expression, confirming a cell autonomous role for insulin signaling in the intestine (**Extended Data Fig. 4C**). Consistent with the non-cell autonomous increase in intestinal DAF-16/FOXO, we observed that neuron-specific knockdown of *egl-3/PC2* also upregulated intestinal MTL-1 (**Fig. 3F**), confirming the neuron-to-intestine signaling. Together, these results show that attenuating neuron-to-intestine peptide signaling, non-cell autonomously activates intestinal DAF-16/FOXO, that may reprogram the intestine to delay its age-related pathologies and improve survival outcomes.

### Neuropeptide signaling acts on intestinal epithelial barrier components in a DAF-16/FOXO-dependent manner

We hypothesized that specific DAF-16/FOXO targets act in the intestine, downstream of neuropeptide signaling, to maintain epithelial barrier homeostasis. We mined a DAF-16/FOXO ChIP-sequencing dataset, in which DAF-16/FOXO was activated in *daf-2(e1370)* reduction-of-function mutants^47^, to identify genes with enriched DAF-16/FOXO recruitment (binding peaks distributed within ±1kb with respect to the transcriptional start site (TSS)^52^). We identified several epithelial barrier components, including the intestine-specific actin; *act-5*, tight-junction components; *claudins, par-3*, and *par-5*, the basolateral component; *let-413*, and septate-junction components; *ajm-1* and *dlg-1*, among the differentially enriched DAF-16/FOXO targets in response to reduced insulin signaling (**Extended Data Fig. 5A**). To test whether these genes act downstream of neuropeptide signaling, we generated transgenic animals expressing the *unc-31/CAPS* hairpin RNAs pan-neuronally in the intestine-specific feeding RNAi transgenics (**Fig. 3G**). This transgenic strain allowed us to test whether an intestine-specific knockdown of a gene of interest can reverse the reduced intestinal leakiness evoked by the neuronal *unc-31/CAPS* knockdown (**Extended Data Fig. 5B**). We first confirmed the effectiveness of this strain, by feeding these animals bacteria expressing dsRNA for *daf-16/FOXO*. We found that knocking down *daf-16/FOXO* in the intestine increased its leakiness in the *unc-31/CAPS* hairpin-RNAi transgenics, confirming its sensitivity to feeding RNAi and a role for intestinal DAF-16/FOXO in promoting intestinal integrity (**Fig. 3H**). Next, we tested whether intestine-specific knockdown of epithelial components, identified as direct DAF-16/FOXO targets, had a role in regulating epithelial barrier function. We found that intestine-specific knockdown of *act-5* (actin) (**Fig. 3I**), *ajm-1* (septate junction component) (**Fig. 3J**), *clc-1* (tight junction component) (**Extended Data Fig. 5C**) or *let-413* (a basolateral component) (**Extended Data Fig. 5D**), significantly suppressed the HypoSmurf phenotype of *unc-31/CAPS* hairpin-RNAi transgenics. Together, these results show that blocking neuropeptide signaling engage key epithelial barrier components to maintain barrier integrity, possibly via a DAF-16/FOXO-dependent transcriptional mechanism.

**Fig. 5.**
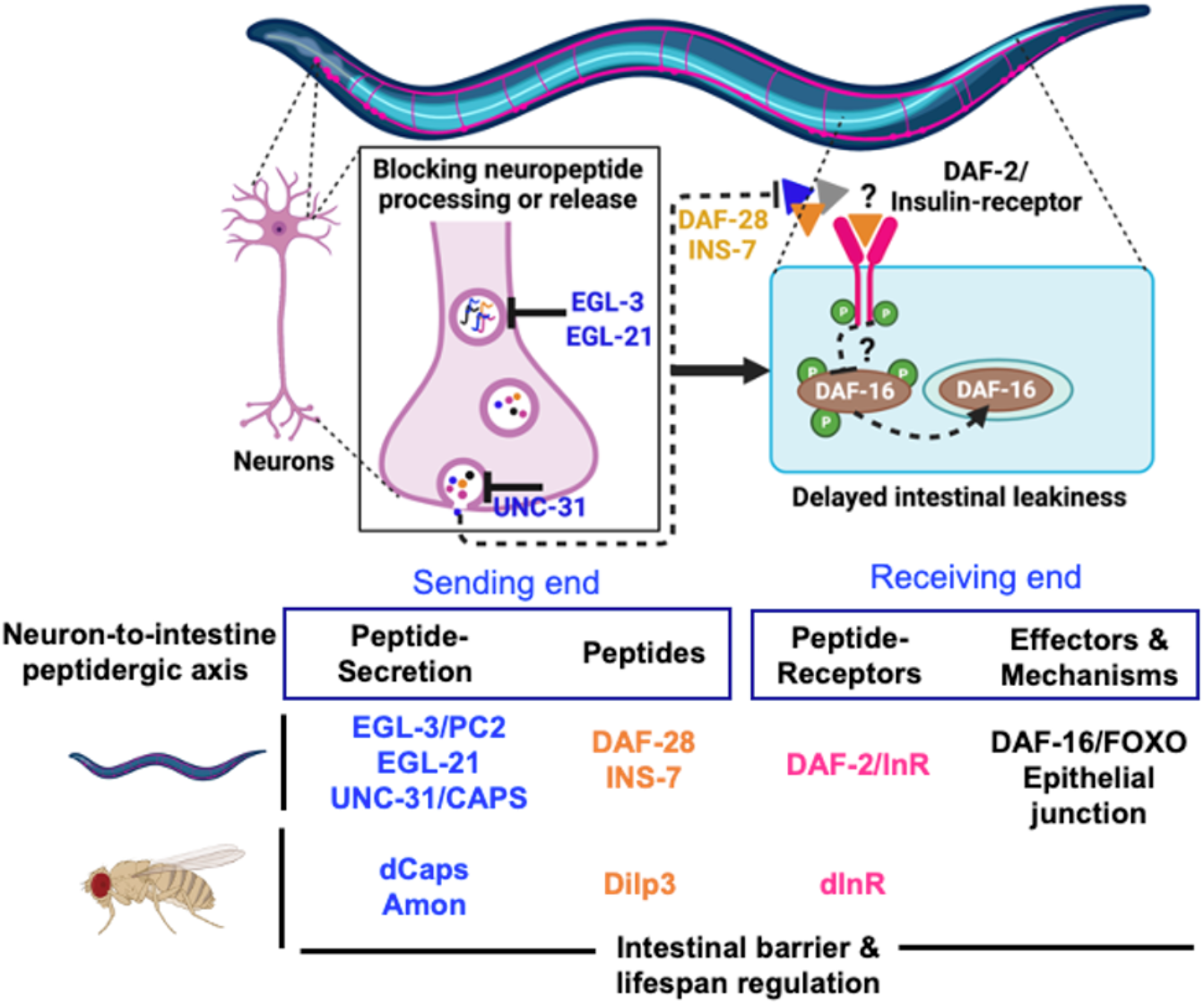
Conserved neuropeptidergic modulation of intestinal integrity in *C. elegans* and *D. melanogaster*. Model showing conserved components in the intestine-modifying neuropeptide signaling pathway. In *C. elegans*, we also show that intestine-modifying neuropeptide signaling impinges upon DAF-16/FOXO and components of the intestinal epithelial barrier. Also see text for details.

### Secretion of DAF-28 peptides correlates with reproductive aging and onset of intestinal aging

As the age-related increase in intestinal leakiness is predominantly post-reproductive (**Extended Data Fig. 6A**), we speculated that the upstream neuropeptide secretion might also be affected by germline activity. Therefore, we turned our focus to quantify the age-related accumulation of GFP-tagged DAF-28 in coelomocytes^53,54^. DAF-28 neuropeptides are expressed in head (ASI, ASJ) and tail (PQR) sensory neurons and the distal intestine^39^ and accumulates in the coelomocytes upon its secretion (**Fig. 4A-B**). Neuropeptide secretion has been previous quantified by monitoring the accumulation of fluorescently tagged peptides in coelomocytes^55^. Although the levels of DAF-28 peptides (measured as GFP fluorescence in head neurons) was comparable during the reproductive phase (**Extended Data Fig. 6B**), secretion of DAF-28 (quantified as the number of GFP-positive coelomocytes) exhibited an age-related surge and decline (**Fig. 4C)**. Interestingly, this DAF-28 secretion profile mirrored the reproductive span (**Fig. 4D**)^56,57^, consistent with its function in promoting reproduction^58^.

When considering the relationships among reproductive aging, age-related DAF-28 secretion, and intestinal aging, we predicted that altering germline function could possibly impact intestinal aging either directly or by altering the secretion of DAF-28 peptides (**Fig. 4E**). To this end, we analyzed germline-less *glp-1(e2144ts)* mutants, which exhibited delayed intestinal leakiness (**Fig. 4F**). Importantly, *glp-1(e2144ts)* animals are longer lived, at least in part due to intestinal DAF-16/FOXO activation^11,59^; however, the inter-tissue signals are not understood. We therefore tested if a neuronal insulin-peptide circuit could be part of the effects on the intestinal barrier function. In support of this notion, we found that neuronal over-expression of *daf-28* was sufficient to hasten the observed delay in intestinal leakiness in germline-less *glp-1(e2144ts)* mutants (**Fig. 4G**). Collectively, our results suggest that the *C. elegans* germline activity may engage neurons to regulate the secretion of intestine-modifying neuropeptides.

## Discussion

We find that neuron-specific knockdown of *CAPS* (the activator protein for peptide secretion) and the peptide-processing enzyme, prohormone convertase 2 (PC2), in young adults reduced age-related intestinal leakiness and extended survival in both *C. elegans* and *D. melanogaster*. Furthermore, using a neuron-specific genetic screen for insulin-like peptides, we identified antagonizing roles in the maintenance of intestinal barrier integrity for two *C. elegans* insulins, *daf-28* and *ins-7*, and three *D. melanogaster dilp2, dilp3*, and *dilp5* insulins. On the receiving end, knocking down *daf-2/InR* insulin receptor or the downstream *age-1/PI3K* cell-autonomously improved intestinal health, and lifespan. In *C. elegans*, we showed that reducing neuropeptide signaling non-cell-autonomously activated intestinal DAF-16/FOXO to potentially upregulate the transcription of genes associated with the intestinal epithelial barrier (*act-5, clc-1, let-413, dlg-1, ajm-1*, and others). Broadly, our results establish a conserved role for neuropeptides in affecting barrier function in both *C. elegans* and *D. melanogaster* intestines (**Fig. 5**)

Intestinal barriers are disrupted with age or in humans with Type 1 diabetes, or neurodegenerative conditions, but its genetic regulation is not well studied^60,61^. We show that neuronal peptides affect intestinal integrity in both nematodes and flies, and identify the proteolytic processing of pro-peptides by pro-protein convertases (PC2)^32^ as a key step in this process. Consistently, both *C. elegans egl-3/PC2* and *D. melanogaster amon/PC2* mutants are deficient in bioactive neuropeptides^36,62^. Moreover, deficits in neuropeptide signaling result in developmental and growth defects in nematodes, flies, and mice^63-66^ and, disease states in humans^67,68^. However, using a neuron-specific knockdown approach we show that knocking down peptide processing in young adults leads to improved intestinal health and increased longevity, in a DAF-16/FOXO-dependent manner. Furthermore, over-expressing *egl-3/PC2* in *C. elegans* neurons worsened intestinal leakiness in aged adults, likely by increasing the level of bioactive peptides, which are then secreted to exacerbate intestinal integrity, likely by inducing insulin signaling. We suggest that EGL-3/PC2-mediated neuropeptide processing is a rate-determining step in intestinal aging.

While genes involved in cytoskeleton organization are conserved DAF-16/FOXO direct targets in nematodes and flies^47,69^, its role in regulating epithelial barriers components is understudied. DAF-16/FOXO has increased promoter occupancy on genes associated with intestinal epithelial junctions, including *act-5, let-413* and *dlg-1*^52^ and we further validated a functional role for these proteins in modifying intestinal integrity, downstream of the neuron-to-intestine peptide signaling. Consistent with our findings in *C. elegans*, dFOXO binding to the *Dlg1* promoter is greatly reduced in aged flies^70^, likely reducing *Dlg1* expression and increasing barrier dysfunction in these animals. Together, we suggest that DAF-16/FOXO maintains the intestinal epithelial barrier function, by upregulating stress response genes like *mtl-1* and, building blocks of epithelial junctions, like *act-5, claudins, let-413, ajm-1, dlg-1* etc. Supporting this notion, previous studies in flies have shown that depleting septate junction components is sufficient to exacerbate barrier function in young flies, similar to the impact of physiological age^71^.

Decline in germline function and reproductive capacity is one of the early aging phenotypes^72^. Nematode DAF-28 and fly Dilp3 are both required for proper germline maturation during development^58,73^, however these peptides become undetectable in post-reproductive adult nematodes and, flies^74^. Collectively, these data suggest an intriguing hypothesis – that germline activity modifies neuronal signaling, which in turn impacts intestinal aging. Consistent with this hypothesis, *C. elegans* adults lacking germline exhibit primitive chemotaxis behavior like those of developing larvae, as germline proliferation dictates animal behavior by altering responsiveness of olfactory neurons^75^. Also, a role for reproductive senescence in modifying neuronal signaling has been observed in multiple species including humans^76-78^. Here, we posit that besides modulating adult behaviors, germline signals govern the secretion of intestine-modifying neuropeptides to alter aging, primarily in the intestine. Consistent with this view, we observed that *C. elegans glp-1* mutants, which lack a germline and are long-lived, displayed improved intestinal integrity, a phenotype that was significantly suppressed by neuronal overexpression of *daf-28*. We suggest that the *C. elegans* germline engages neurons, possibly in an age-dependent manner, to produce intestine-modifying neuropeptides that becomes a trade-off for subsequent intestinal aging. In the long-term, these studies can provide scientific frameworks to establish the etiology of intestinal aging in mammals and altered brain-gut communications during age-related neuropathies.

## Materials and Methods

### Bacterial strains

*Escherichia coli* OP50 was obtained from Dr. Cynthia Kenyon (formerly at UCSF), grown at 37ºC in Luria Bertani (LB) broth. For all RNAi experiments, HT115 *E. coli* were cultured under carbenicillin selection. All HT115 bacteria were grown for 6-8 hours at 37ºC in Luria Bertani (LB) broth, cultures were shaken at 220 rpm.

### C. elegans strains and husbandry

*C. elegans* were maintained on nematode growth media plates seeded with *E. coli* OP50 at 20°C^79^. For RNAi experiments, animals were grown on HT115 bacteria from day 1 of gravid adulthood (unless otherwise indicated). All *C. elegans* strains used in this study are listed in **Supplementary Table 5**. Bristol N2, both from Caenorhabditis Genetics Center (N2-CGC) and the Kenyon lab (N2-CK) were used as the wild-type animal (as indicated). For experiments with temperature-sensitive *glp-1(e2144ts)* mutants, both wild-type (N2-CK) and *glp-1(e2144ts)* stage 4 larva (L4) were shifted to 25ºC for egg-laying and until these progenies developed to day 1 of adulthood.

### Drosophila lines and husbandry

All fly stocks were maintained at 25ºC or room temperature on standard fly food containing cornmeal-molasses agar media. Lines obtained from Bloomington *Drosophila* Stock Center (BDSC, Bloomington, IN): *w1118* (#5905), elav^C155^; TubGal80 (#67058), UAS-RNAi-*cadps* (# 31984), UAS-RNAi-*Amon* (#28583), UAS-RNAi-*dilp2* (#32475), UAS-RNAi-*dilp3* (#33681), UAS-RNAi-*dilp5* (#33683), and UAS-RNAi-*InR* (#51518), NP1-Gal4, TubGal80/*cyo*^80^ and UAS-GFP/MKRS were available in the Bier lab.

## Method details

### RNA interference in C. elegans

Tissue-specific RNAi in the *C. elegans* model was implemented by either using feeding-RNAi method or hairpin-mediated RNAi^81^. Tissue-specific expression of the *sid-1* cDNA (codes for membrane-associated, double-stranded RNA transporter), in a *sid-1(qt9)* null mutant, limits the RNAi activity to a particular tissue^82^. Neuron-specific or intestine-specific RNA interference (RNAi) transgenics are generated by expressing the *sid-1* cDNA using a pan-neuronal *rgef-1* promoter (Y. Yang, C. Kumsta and M. Hansen, manuscript in preparation)^31^ or intestinal *vha-6p* promoter^83^, in a *sid-1(qt9)* null mutant. For hairpin-mediated RNAi, *unc-31/CAPS* was knocked down by pan-neuronally expressing *unc-31* hairpin RNAs under *h20* promoter^81^. For feeding-RNAi method, gene knockdown was implemented by feeding with *E. coli* HT115 bacterial clones expressing double-stranded RNA (dsRNA) targeting the gene of interest. *unc-31/CAPS* RNAi clone was generated for this study, other RNAi bacterial clones were obtained from the Ahringer RNAi library^84^ or the Vidal RNAi library^85^, and *daf-2, age-1* and *daf-16* RNAi clones were provided by Dr. Andrew Dillin (UC Berkeley), the empty vector (L4440) for RNAi controls and *daf-28* RNAi clone (from Vidal library) were provided by Drs. Andrew Fire (Stanford University) and Arnab Mukhopadhyay (National Institute of Immunology), respectively. All bacterial clones used in this study were verified by sequencing (**Supplementary Table 6**). For feeding-RNAi experiments, HT115 bacterial clones were grown in Luria-Bertani (LB) liquid culture medium containing 0.1 mg ml^−1^ Carbenicillin (Carb; BioPioneer) for 6-8 hours, 140 μl aliquots of bacterial suspension were seeded onto 6 cm Petri plates with NGM containing 0.1 mg ml^−1^ Carbenicillin (NGM-Carb plates). Bacteria were allowed to grow for 1–2 days and induced for dsRNA expression by adding 140 μl of an isopropyl-[3-D-thiogalactoside solution containing 0.1M IPTG (Promega/Gold Biotechnology) and 500 mg ml^−1^ Carbenicillin added directly onto the bacterial lawn. For adult-only RNAi experiments, eggs laid by day 1 gravid adults were collected onto NGM plates seeded with OP50 bacteria for hatching and development till day 1 of adulthood. Day 1 gravid adults were transferred to NGM-Carb plates seeded with bacteria expressing control or dsRNA for indicated genes.

### Construction of unc-31 RNAi clone

1160-base-pair of *unc-31* gDNA was amplified from wild-type genomic DNA using the forward primer 5’-CCACCGGTCTTCACCGCATTCGACAGTT-3’ and the reverse primer 5’-CTAGCTAGCCAA GCTGTAATCGTCGACTGA-3’ using NEB Taq Polymerase, cloned into AgeI and NheI restriction sites of pL4440 vector and transformed into *E. coli* HT115(DE3) cells.

### RNA interference in Drosophila

Tissue-specific RNAi in the *Drosophila* model was implemented using the binary GAL4-UAS (Upstream Activating Sequences) system^86^. Tissue-specific Gal4 drivers limited the tissue-specific expression of RNAi^87^ and timing of GAL4 activity was limited by a temperature sensitive form of GAL80 (*tubulin*-GAL80ts), which inhibits GAL4 function below 25°C. In this study, for adult-only neuron-specific knockdown of *cadps, amon, dilp2, dilp3* or *dilp5*, neuron-specific Gal4: elav^C155^; TubGal80 flies^88^, were crossed to UAS-RNAi-*cadps*, UAS-RNAi-*amon*, UAS-RNAi-*dilp2*, UAS-RNAi-*dilp3 or* UAS-RNAi-*dilp5* flies, respectively and, 1-2 days old female F1 flies of desired genotype were shifted to 29ºC for 10 days (or indicated otherwise) for neuron-specific GAL4 activation. Similarly, for adult-only gut-specific knockdown of *InR*, NP1-Gal4, TubGal80/*cyo* flies were crossed to UAS-RNAi-*InR* flies, and 3-5 days old female F1 flies of desired genotype were shifted to 29ºC for 10 days (or indicated otherwise) for gut-specific GAL4 activation.

### Lifespan assay in C. elegans

Lifespan experiments were performed at 20ºC with ∼100 animals per condition, with 12–15 animals on each 6 cm NGM-Carb petri plates seeded with bacteria expressing empty control vector or dsRNA against the indicated gene. Feeding-RNAi was initiated on day 1 of adulthood and maintained throughout the lifespan analysis unless indicated otherwise. The L4 larval stage was recorded as day 0 of the lifespan and animals were transferred at least every other day to new NGM plates throughout the reproductive period. Animals were censored if they ruptured, escaped, desiccated on the edge of the plate, or suffered from internal hatching or contamination. OASIS2 software was used for statistical analyses and *P-values* were calculated using the log-rank (Mantel– Cox) test, a nonparametric test used to compare two survival functions through overall lifespan data^89^. See Supplementary **Tables S2** for a summary of all lifespan experiments.

### Lifespan assay in D. melanogaster

Lifespan experiments were performed at 25ºC with more than 50 animals per condition (protocol adapted from^90^. For neuron-specific RNAi experiments, 30-40 *elav*^*C155*^; TubGal80 virgin females were crossed to 15-25 males of desired UAS-RNAi line, or *w1118* (wild type) and maintained at 25°C. Similarly, for gut-specific RNAi experiments, 30-40 NP1-Gal4, TubGal80/*cyo* virgins were crossed to 15-25 males of UAS-RNAi-*InR* or *w1118* and maintained at 25°C. F1 flies were collected and maintained at 25°C for a day, anaesthetized by CO_2_ for sorting by sex and transferred to vials (10-15 flies/vial). Female flies were used for all lifespan experiments if not indicated otherwise. 1– 2-day old female flies were shifted to 29°C for 10 days (or indicated otherwise) for tissue-specific RNAi induction and then maintained at 25°C throughout lifespan. The dead flies were scored every alternate day and flies were transferred to fresh food vials twice a week (no CO_2_ exposure). OASIS2 software^89^ was used for statistical analyses and *P values* were calculated using the log-rank (Mantel–Cox) test. See **Supplementary Table 3** for a summary of all lifespan experiments.

### Smurf and FluoroSmurf assay in C. elegans

For Smurf assays, animals were aged on NGM-Carb plates supplemented with 120 μl of 5-Fluoro-2′-deoxyuridine solution containing 10 μl of 0.4M FUdR (Sigma-Aldrich, F0503) to prevent hatching of progenies and maintain synchronous nematode populations (Mitchell *et al*., 1979). For Smurf assay, day 15 old nematodes (unless otherwise mentioned) were fed 250 μl of 5.0% wt/vol stock of blue food dye (Spectrum FD&C Blue #1 PD110) dissolved in overnight grown culture of OP50 bacteria^21^, in 96-well flat bottom plate. Following 3 hours of dye feeding at 20°C, animals were visualized under a microscope and classified into either Smurfs or non-Smurfs based on blue dye retention, and % Smurf was calculated. Animals with blue dye only in their germline were excluded from total population^21^. To quantify intestinal leakiness, we implemented modified FluoroSmurf assay by combining fluorescein with the blue dye. For these FluoroSmurf assays, worms were fed 125μl each of blue food dye and 2.5% wt/vol stock of Fluorescein sodium salt (Sigma-Aldrich, F6377), each dissolved in an overnight-grown culture of OP50 bacteria. Following 3 hours of dye feeding at 20°C, animals were first visualized under a microscope and classified into either Smurfs or non-Smurfs based on blue dye retention to calculate % Smurf. Next, animals were anaesthetized on NGM plates without food with M9 buffer containing 150mM NaN_3_, classified into Smurfs on non-Smurfs based on blue dye localization and imaged with a Leica DFC310 FX camera to visualize and measure fluorescein dye accumulation. Exposure time was kept constant within an experiment (3ms, 5ms, or 20ms), but varied between experiments and therefore absolute fluorescence of day 15 control animals also varied (higher absolute fluorescence at higher exposure). Fluorescein intensity was measured by outlining the entire animal using polygon tool of the FIJI/Image J software and, reported in an arbitrary unit relative to wild type/control. Age-related intestinal leakiness was monitored in cross-sectional Smurf assays, such that day 5, day 10, and day15 old nematodes populations were assayed on the same day.

### Smurf assay in D. melanogaster

Longitudinal studies were conducted to monitor both lifespan and intestinal leakiness in aging fly population. Smurf assay was adapted from^91^. Briefly, 4 g of blue dye (Spectrum FD&C Blue #1 PD110) was dissolved in 10 ml distilled water. 7.8 ml of this stock was added to 100 ml of *Drosophila* food and distributed into 3 ml per vial. Care was taken to avoid any condensation and vials were set aside at room temperature and used the next day. On the day of Smurf assay, flies were transferred to blue-food and maintained at 25°C. After 24 hr of dye-feeding, flies were anaesthetized by CO_2_ to monitor the blue dye localization under a microscope. As in nematodes, flies were classified as ‘Smurfs’ when the blue dye leaked into their body cavity. Following Smurf assays at multiple age-points, live flies were transferred back to fresh food vials for further lifespan analysis.

### C. elegans motility assay

Thrashing rate was monitored at different age-points as indicated in the corresponding figures by counting the number of body bends per 30 seconds of randomly picked *C. elegans* swimming on NGM plate in a drop of M9 buffer. Movement of head was monitored under a stereo microscope, counted as one body bend when animal moved its anterior body from one side to another. Statistical comparisons were performed using two-way ANOVA, with Tukey’s multiple comparisons test, on data pooled from two independent experiments (n=2), taking 10 animals per condition in each set.

### C. elegans pharyngeal pumping assay

At different age-points as indicated in the corresponding figures, the number of contractions of the pharyngeal terminal bulb were counted over 30 seconds. Animals were randomly picked and observed under a compound microscope, and a contraction was defined as posterior movement of the grinder in the pharyngeal terminal bulb. Statistical comparisons were performed using two-way ANOVA, with Tukey’s multiple comparisons test, on data pooled from two independent experiments (n=2), taking 10 animals per condition in each set.

### C. elegans defecation and gastric motility assay

Posterior body was monitored for contraction of anal muscles (expulsion step, following posterior and anterior body-wall muscle contractions) under a compound microscope and length of defecation cycle was recorded as time in seconds, between two contractions of anal muscles ^92^. In a new approach to measure gastric motility, animals were first fed OP50 bacteria mixed with blue dye for an hour and the presence of blue dye in the intestinal lumen was confirmed. Animals were then fed fresh OP50, and dye expulsion was monitored, visualized as loss of blue dye from the intestinal lumen. Using this approach, gastric motility of an animal was reported as a ratio of time taken for dye expulsion (in seconds) and length of defecation cycle (in seconds). Statistical comparisons were performed by two-tailed *t* test for unpaired samples, data pooled from three independent experiments (n=3), taking 10 animals per condition in each set.

### C. elegans reproductive span

Animals were first synchronized by egg laying and raised on nematode growth media (NGM) plates seeded with *E. coli* OP50 at 20°C, until L4 stage. Then, 10 randomly picked L4 larvae were singly plated onto NGM plates with *E. coli* OP50. Adult worms were transferred every 24 hours onto fresh OP50 plates till the end of self-fertile period. To determine the number of viable progenies laid each day, number of L1 larvae were counted 24 hours after adult worms were moved, and 48 hours later for counting L1s that developed into adult worms. The reproductive span was reported as number of viable adult progenies generated each day of the self-fertile period.

### Coelomocyte assay to monitor peptide secretion in C. elegans

Coelomocyte assay was performed to monitor age-related DAF-28 peptide secretion^53^. To determine the age-dependent secretion of DAF-28 insulin-like peptide, *daf-28p::daf-28::gfp* transgenics were aged simultaneously, such that day 1 to day 8 old animals were anaesthetized (150mM NaN_3_) and imaged on the same day using a Leica DFC310 FX camera. Age-dependent secretion of DAF-28 insulin-like peptide was reported as the average number of GFP-positive coelomocytes per animal (at least 10 animals were monitored per age-point). To estimate the pre-secreted DAF-28 peptides synthesis, fluorescence was quantified from the ASI/ASJ head neurons.

### Generation of transgenic animals

#### rgef-1p::egl-3c cDNA::pSM::mCherry

The 1584 bp of *egl-3* cDNA was amplified from wild-type cDNA using a forward primer (5’ ctaGCTAGCATGTCTCCATCGGATCCAC 3’) and reverse primer (5’ cggGGTACCTTAGTGGCTGCGTTTGTGG 3’) using Q5^®^ High-Fidelity DNA Polymerase system (New England Biolabs, USA) and cloned into *rgef-1p::pSM::mCherry* vector using *NheI* and *KpnI* restriction sites. The recombinant plasmid was microinjected at a concentration of 100ng/μl into the syncytial gonad of wild-type N2 (CGC) worms.

#### rgef-1p::daf-28(SM)::pSM::mCherry

The 700 bp of *daf-28* gDNA was amplified from *daf-28p::daf-28::mCherry* plasmid provided by Dr. Shohei Mitani (Tokyo Women’s Medical University) using a forward primer (5’ ctaGCTAGCATGAACTGCAAGCTCATCGC 3’) and reverse primer (5’ cggGGTACCTTAAAGAAGCAAACGTGGGCA 3’) using Q5^®^ High-Fidelity DNA Polymerase system (New England Biolabs, USA) and cloned into *rgef-1p::pSM::mCherry* vector using *NheI* and *KpnI* restriction sites. The recombinant plasmid was microinjected at a concentration of 80ng/μl into the syncytial gonad of wild-type N2 (CGC) worms.

#### rgef-1p::ins-7(2b)::pSM::mCherry

The 764 bp of *ins-7(2b)* gDNA was amplified from wild-type genomic DNA using a forward primer (5’ ctaGCTAGCATGTATAAAGTACATTATTTTC 3’) and reverse primer (5’ cggGGTACCTTAAGGACAGCACTGTTTT 3’) using Q5^®^ High-Fidelity DNA Polymerase system (New England Biolabs, USA) and cloned into *rgef-1p::pSM::mCherry* vector using *NheI* and *KpnI* restriction sites. The recombinant plasmid was microinjected at a concentration of 80ng/μl into the syncytial gonad of wild-type N2 (CGC) worms.

## Data availability

The authors declare that all data supporting the findings of this study are available within this article. ChIP-Atlas database^52^ was used to determine DAF-16 targets using data from DAF-16 ChIP sequencing study^47^ available from the Gene Expression Omnibus (GSE63865 and GSM1558630).

## Code availability

None were used in this study

## Acknowledgements

We thank Dr. Yongzhi Yang, Chung-Chih Liu, Andrew Davis, and Linnea Adams for technical assistance. We thank Drs. Andrew Dillin, Andrew Fire, Arnab Mukhopadhyay, Cynthia Kenyon, Mei Zhen, Shohei Mitani and Supriya Srinivasan, for kindly providing strains and reagents. We are also grateful to Drs. Rolf Bodmer, Karen Ocorr, and Annabel Guichard and, members of the Chalasani and Hansen labs for critical comments and insights. Some of the nematode strains used in this work were provided by the Caenorhabditis Genetics Center (University of Minnesota), which is supported by the NIH–Office of Research Infra-structure Program (P40 OD010440).

## Funding

National Institutes of Health grant MH096881 (SHC)

National Institutes of Health grant AG038664 (MH)

Breakthrough in Gerontology Award from the American Federation for Aging Research (MH)

Postdoctoral fellowship from the Paul F. Glenn Center for Biology of Aging Research at the Salk Institute (AS)

## Author contributions

Conceptualization: AS, MH, SHC

Methodology: AS, BK, KCR, CK

Investigation: AS, BK

Visualization: AS, BK

Funding acquisition: MH, SHC

Project administration: AS, MH, SHC

Supervision: EB, MH, SHC

Writing – original draft: AS, SHC

Writing – review & editing: AS, SHC, MH, CK, BK, EB, KCR

## Competing interests

Authors declare that they have no competing interests.

## Data and materials availability

All data are available in the main text or the supplementary materials. Further information and requests for resources and reagents should be directed to and will be fulfilled by the Lead Contact, Sreekanth H Chalasani. *C. elegans* strains and RNAi constructs generated in this study are available upon request from the Lead Contact with a completed Materials Transfer Agreement.

## Notes

### Competing Interest Statement

The authors have declared no competing interest.

## References

1 Arrojo, E. D. R. et al. Age Mosaicism across Multiple Scales in Adult Tissues. Cell Metab 30, 343–351 e343, doi:10.1016/j.cmet.2019.05.010 (2019).

2 Miller, H. A., Dean, E. S., Pletcher, S. D. & Leiser, S. F. Cell non-autonomous regulation of health and longevity. eLife 9, e62659, doi:10.7554/eLife.62659 (2020).

3 Katzenberger, R. J. et al. Death following traumatic brain injury in Drosophila is associated with intestinal barrier dysfunction. eLife 4, doi:10.7554/eLife.04790 (2015).

4 Ma, E. L. et al. Bidirectional brain-gut interactions and chronic pathological changes after traumatic brain injury in mice. Brain Behav Immun 66, 56–69, doi:10.1016/j.bbi.2017.06.018 (2017).

5 Winge, K., Rasmussen, D. & Werdelin, L. M. Constipation in neurological diseases. J Neurol Neurosurg Psychiatry 74, 13–19 (2003).

6 Katsimpardi, L. et al. Systemic GDF11 stimulates the secretion of adiponectin and induces a calorie restriction-like phenotype in aged mice. Aging Cell 19, e13038, doi:10.1111/acel.13038 (2020).

7 Castellano, J. M. Blood-Based Therapies to Combat Aging. Gerontology 65, 84–89, doi:10.1159/000492573 (2019).

8 Fan, X., Gaur, U., Sun, L., Yang, D. & Yang, M. The Growth Differentiation Factor 11 (GDF11) and Myostatin (MSTN) in tissue specific aging. Mech Ageing Dev 164, 108–112, doi:10.1016/j.mad.2017.04.009 (2017).

9 Glatigny, M. et al. Autophagy Is Required for Memory Formation and Reverses Age-Related Memory Decline. Curr Biol 29, 435–448 e438, doi:10.1016/j.cub.2018.12.021 (2019).

10 Apfeld, J. & Kenyon, C. Cell nonautonomy of C. elegans daf-2 function in the regulation of diapause and life span. Cell 95, 199–210, doi:S0092-8674(00)81751-1 [pii] (1998).

11 Libina, N., Berman, J. R. & Kenyon, C. Tissue-specific activities of C. elegans DAF-16 in the regulation of lifespan. Cell 115, 489–502, doi:S0092867403008894 [pii] (2003).

12 Wolkow, C. A., Kimura, K. D., Lee, M. S. & Ruvkun, G. Regulation of C. elegans life-span by insulinlike signaling in the nervous system. Science 290, 147–150 (2000).

13 Masse, I., Molin, L., Billaud, M. & Solari, F. Lifespan and dauer regulation by tissue-specific activities of Caenorhabditis elegans DAF-18. Dev Biol 286, 91–101, doi:10.1016/j.ydbio.2005.07.010 (2005).

14 Ailion, M., Inoue, T., Weaver, C. I., Holdcraft, R. W. & Thomas, J. H. Neurosecretory control of aging in Caenorhabditis elegans. Proc Natl Acad Sci U S A 96, 7394–7397 (1999).

15 Leinwand, S. G. et al. Circuit mechanisms encoding odors and driving aging-associated behavioral declines in Caenorhabditis elegans. eLife 4, e10181, doi:10.7554/eLife.10181 (2015).

16 Liu, J. et al. Functional aging in the nervous system contributes to age-dependent motor activity decline in C. elegans. Cell Metab 18, 392–402, doi:10.1016/j.cmet.2013.08.007 (2013).

17 Li, G. et al. Genetic and pharmacological interventions in the aging motor nervous system slow motor aging and extend life span in C. elegans. Science advances 5, eaau5041 (2019).

18 Ching, T.-T. et al. Short-term enhancement of motor neuron synaptic exocytosis during early aging extends lifespan in Caenorhabditis elegans. Experimental Biology and Medicine 245, 1552–1559 (2020).

19 Zullo, J. M. et al. Regulation of lifespan by neural excitation and REST. Nature 574, 359–364, doi:10.1038/s41586-019-1647-8 (2019).

20 Carrano, A. C., Dillin, A. & Hunter, T. A Krüppel-like factor downstream of the E3 ligase WWP-1 mediates dietary-restriction-induced longevity in Caenorhabditis elegans. Nature communications 5, 1–9 (2014).

21 Gelino, S. et al. Intestinal Autophagy Improves Healthspan and Longevity in C. elegans during Dietary Restriction. PLoS Genet 12, e1006135, doi:10.1371/journal.pgen.1006135 (2016).

22 Rera, M., Azizi, M. J. & Walker, D. W. Organ-specific mediation of lifespan extension: more than a gut feeling? Ageing Res Rev 12, 436–444, doi:10.1016/j.arr.2012.05.003 (2013).

23 Dambroise, E. et al. Two phases of aging separated by the Smurf transition as a public path to death. Sci Rep 6, 23523, doi:10.1038/srep23523 (2016).

24 Meier, J. & Sturm, A. The intestinal epithelial barrier: does it become impaired with age? Dig Dis 27, 240–245, doi:10.1159/000228556 (2009).

25 Rera, M., Clark, R. I. & Walker, D. W. Intestinal barrier dysfunction links metabolic and inflammatory markers of aging to death in Drosophila. Proceedings of the National Academy of Sciences 109, 21528–21533 (2012).

26 Buchon, N., Broderick, N. A. & Lemaitre, B. Gut homeostasis in a microbial world: insights from Drosophila melanogaster. Nature Reviews Microbiology 11, 615–626 (2013).

27 Miguel-Aliaga, I., Jasper, H. & Lemaitre, B. Anatomy and physiology of the digestive tract of Drosophila melanogaster. Genetics 210, 357–396 (2018).

28 Richmond, J. E., Davis, W. S. & Jorgensen, E. M. UNC-13 is required for synaptic vesicle fusion in C. elegans. Nat Neurosci 2, 959–964 (1999).

29 Nonet, M. L., Saifee, O., Zhao, H., Rand, J. B. & Wei, L. Synaptic transmission deficits in Caenorhabditis elegans synaptobrevin mutants. J Neurosci 18, 70–80 (1998).

30 Speese, S. et al. UNC-31 (CAPS) is required for dense-core vesicle but not synaptic vesicle exocytosis in Caenorhabditis elegans. J Neurosci 27, 6150–6162 (2007).

31 Calixto, A., Chelur, D., Topalidou, I., Chen, X. & Chalfie, M. Enhanced neuronal RNAi in C. elegans using SID-1. Nat Methods 7, 554–559, doi:nmeth.1463 [pii] 10.1038/nmeth.1463 (2010).

32 Li, C. & Kim, K. Neuropeptides. WormBook, 1–36 (2008).

33 Lindberg, I., Tu, B., Muller, L. & Dickerson, I. M. Cloning and functional analysis of C. elegans 7B2. DNA Cell Biol 17, 727–734, doi:10.1089/dna.1998.17.727 (1998).

34 Van Bael, S. et al. Mass spectrometric evidence for neuropeptide-amidating enzymes in Caenorhabditis elegans. J Biol Chem 293, 6052–6063, doi:10.1074/jbc.RA117.000731 (2018).

35 Renden, R. et al. Drosophila CAPS is an essential gene that regulates dense-core vesicle release and synaptic vesicle fusion. Neuron 31, 421–437 (2001).

36 Wegener, C., Herbert, H., Kahnt, J., Bender, M. & Rhea, J. M. Deficiency of prohormone convertase dPC2 (AMONTILLADO) results in impaired production of bioactive neuropeptide hormones in Drosophila. J Neurochem 118, 581–595, doi:10.1111/j.1471-4159.2010.07130.x (2011).

37 Taylor, S. R. et al. Molecular topography of an entire nervous system. Cell 184, 4329–4347 e4323, doi:10.1016/j.cell.2021.06.023 (2021).

38 Fernandes de Abreu, D.A. et al. An insulin-to-insulin regulatory network orchestrates phenotypic specificity in development and physiology. PLoS Genet 10, e1004225, doi:10.1371/journal.pgen.1004225 (2014).

39 Li, W., Kennedy, S. G. & Ruvkun, G. daf-28 encodes a C. elegans insulin superfamily member that is regulated by environmental cues and acts in the DAF-2 signaling pathway. Genes Dev 17, 844–858 (2003).

40 Pierce, S. B. et al. Regulation of DAF-2 receptor signaling by human insulin and ins-1, a member of the unusually large and diverse C. elegans insulin gene family. Genes Dev 15, 672–686 (2001).

41 Zheng, S. et al. A functional study of all 40 Caenorhabditis elegans insulin-like peptides. Journal of Biological Chemistry 293, 16912–16922 (2018).

42 Nässel, D. R., Kubrak, O. A., Liu, Y., Luo, J. & Lushchak, O. V. Factors that regulate insulin producing cells and their output in Drosophila. Frontiers in physiology 4, 252 (2013).

43 Nässel, D. R. & Broeck, J. V. Insulin/IGF signaling in Drosophila and other insects: factors that regulate production, release and post-release action of the insulin-like peptides. Cellular and Molecular Life Sciences 73, 271–290 (2016).

44 Kimura, K. D., Tissenbaum, H. A., Liu, Y. & Ruvkun, G. daf-2, an insulin receptor-like gene that regulates longevity and diapause in Caenorhabditis elegans. Science 277, 942–946 (1997).

45 Morris, J. Z., Tissenbaum, H. A. & Ruvkun, G. A phosphatidylinositol-3-OH kinase family member regulating longevity and diapause in Caenorhabditis elegans. Nature 382, 536–539, doi:10.1038/382536a0 (1996).

46 Henderson, S. T. & Johnson, T. E. daf-16 integrates developmental and environmental inputs to mediate aging in the nematode Caenorhabditis elegans. Curr Biol 11, 1975–1980, doi:S0960-9822(01)00594-2 [pii] (2001).

47 Kumar, N. et al. Genome-wide endogenous DAF-16/FOXO recruitment dynamics during lowered insulin signalling in C. elegans. Oncotarget 6, 41418–41433, doi:10.18632/oncotarget.6282 (2015).

48 Oh, S. W. et al. Identification of direct DAF-16 targets controlling longevity, metabolism and diapause by chromatin immunoprecipitation. Nat Genet 38, 251–257, doi:10.1038/ng1723 (2006).

49 Kaletsky, R. et al. The C. elegans adult neuronal IIS/FOXO transcriptome reveals adult phenotype regulators. Nature 529, 92–96, doi:10.1038/nature16483 (2016).

50 Murphy, C. T. et al. Genes that act downstream of DAF-16 to influence the lifespan of Caenorhabditis elegans. Nature 424, 277–283, doi:10.1038/nature01789nature01789 [pii] (2003).

51 Yamawaki, T. M., Arantes-Oliveira, N., Berman, J. R., Zhang, P. & Kenyon, C. Distinct activities of the germline and somatic reproductive tissues in the regulation of Caenorhabditis elegans’ longevity. Genetics 178, 513–526, doi:178/1/513 [pii] 10.1534/genetics.107.083253 (2008).

52 Oki, S. et al. ChIP-Atlas: a data-mining suite powered by full integration of public ChIP-seq data. EMBO Rep 19, doi:10.15252/embr.201846255 (2018).

53 Ch’ng, Q., Sieburth, D. & Kaplan, J. M. Profiling synaptic proteins identifies regulators of insulin secretion and lifespan. PLoS Genet 4, e1000283 (2008).

54 Lee, B. H., Liu, J., Wong, D., Srinivasan, S. & Ashrafi, K. Hyperactive neuroendocrine secretion causes size, feeding, and metabolic defects of C. elegans Bardet-Biedl syndrome mutants. PLoS Biol 9, e1001219, doi:10.1371/journal.pbio.1001219 (2011).

55 Sieburth, D., Madison, J. M. & Kaplan, J. M. PKC-1 regulates secretion of neuropeptides. Nat Neurosci 10, 49–57 (2007).

56 Klass, M. R. Aging in the nematode Caenorhabditis elegans: major biological and environmental factors influencing life span. Mech Ageing Dev 6, 413–429, doi:10.1016/0047-6374(77)90043-4 (1977).

57 Scharf, A., Pohl, F., Egan, B. M., Kocsisova, Z. & Kornfeld, K. Reproductive Aging in Caenorhabditis elegans: From Molecules to Ecology. Front Cell Dev Biol 9, 718522, doi:10.3389/fcell.2021.718522 (2021).

58 Malone, E. A., Inoue, T. & Thomas, J. H. Genetic analysis of the roles of daf-28 and age-1 in regulating Caenorhabditis elegans dauer formation. Genetics 143, 1193–1205 (1996).

59 Berman, J. R. & Kenyon, C. Germ-cell loss extends C. elegans life span through regulation of DAF-16 by kri-1 and lipophilic-hormone signaling. Cell 124, 1055–1068, doi:10.1016/j.cell.2006.01.039 (2006).

60 Saffrey, M. J. Aging of the mammalian gastrointestinal tract: a complex organ system. Age (Dordr) 36, 9603, doi:10.1007/s11357-013-9603-2 (2014).

61 Fan, X. et al. Rapamycin preserves gut homeostasis during Drosophila aging. Oncotarget 6, 35274–35283, doi:10.18632/oncotarget.5895 (2015).

62 Husson, S. J., Clynen, E., Baggerman, G., Janssen, T. & Schoofs, L. Defective processing of neuropeptide precursors in Caenorhabditis elegans lacking proprotein convertase 2 (KPC-2/EGL-3): mutant analysis by mass spectrometry. J Neurochem 98, 1999–2012, doi:JNC4014 [pii] 10.1111/j.1471-4159.2006.04014.x (2006).

63 Furuta, M. et al. Defective prohormone processing and altered pancreatic islet morphology in mice lacking active SPC2. Proc Natl Acad Sci U S A 94, 6646–6651, doi:10.1073/pnas.94.13.6646 (1997).

64 Rayburn, L. Y. et al. amontillado, the Drosophila homolog of the prohormone processing protease PC2, is required during embryogenesis and early larval development. Genetics 163, 227–237, doi:10.1093/genetics/163.1.227 (2003).

65 Siekhaus, D. E. & Fuller, R. S. A role for amontillado, the Drosophila homolog of the neuropeptide precursor processing protease PC2, in triggering hatching behavior. J Neurosci 19, 6942–6954 (1999).

66 Trent, C., Tsuing, N. & Horvitz, H. R. Egg-laying defective mutants of the nematode Caenorhabditis elegans. Genetics 104, 619–647 (1983).

67 Klein-Szanto, A. J. & Bassi, D. E. Proprotein convertase inhibition: Paralyzing the cell’s master switches. Biochem Pharmacol 140, 8–15, doi:10.1016/j.bcp.2017.04.027 (2017).

68 Kovac, S., Shulkes, A. & Baldwin, G. S. Peptide processing and biology in human disease. Curr Opin Endocrinol Diabetes Obes 16, 79–85, doi:10.1097/MED.0b013e3283202555 (2009).

69 Alic, N. et al. Genome-wide dFOXO targets and topology of the transcriptomic response to stress and insulin signalling. Mol Syst Biol 7, 502, doi:10.1038/msb.2011.36 (2011).

70 Birnbaum, A., Wu, X., Tatar, M., Liu, N. & Bai, H. Age-Dependent Changes in Transcription Factor FOXO Targeting in Female Drosophila. Frontiers in genetics 10, 312, doi:10.3389/fgene.2019.00312 (2019).

71 Resnik-Docampo, M. et al. Tricellular junctions regulate intestinal stem cell behaviour to maintain homeostasis. Nature cell biology 19, 52–59 (2017).

72 Quesada-Candela, C., Loose, J., Ghazi, A. & Yanowitz, J. L. Molecular basis of reproductive senescence: insights from model organisms. J Assist Reprod Genet 38, 17–32, doi:10.1007/s10815-020-01959-4 (2021).

73 Dhiman, N. et al. Drosophila Mon1 constitutes a novel node in the brain-gonad axis that is essential for female germline maturation. Development 146, doi:10.1242/dev.166504 (2019).

74 Tanabe, K., Itoh, M. & Tonoki, A. Age-related changes in insulin-like signaling lead to intermediate-term memory impairment in Drosophila. Cell reports 18, 1598–1605 (2017).

75 Fujiwara, M., Aoyama, I., Hino, T., Teramoto, T. & Ishihara, T. Gonadal Maturation Changes Chemotaxis Behavior and Neural Processing in the Olfactory Circuit of Caenorhabditis elegans. Curr Biol 26, 1522–1531, doi:10.1016/j.cub.2016.04.058 (2016).

76 Kermath, B. A. & Gore, A. C. Neuroendocrine control of the transition to reproductive senescence: lessons learned from the female rodent model. Neuroendocrinology 96, 1–12 (2012).

77 Wang, L. et al. Gonadotropin-releasing hormone receptor system: modulatory role in aging and neurodegeneration. CNS & Neurological Disorders-Drug Targets (Formerly Current Drug Targets-CNS & Neurological Disorders) 9, 651–660 (2010).

78 Mosconi, L. et al. Menopause impacts human brain structure, connectivity, energy metabolism, and amyloid-beta deposition. Scientific reports 11, 1–16 (2021).

79 Brenner, S. The genetics of Caenorhabditis elegans. Genetics 77, 71–94 (1974).

80 Jiang, H. et al. Cytokine/Jak/Stat signaling mediates regeneration and homeostasis in the Drosophila midgut. Cell 137, 1343–1355, doi:10.1016/j.cell.2009.05.014 (2009).

81 Esposito, G., Di Schiavi, E., Bergamasco, C. & Bazzicalupo, P. Efficient and cell specific knock-down of gene function in targeted C. elegans neurons. Gene 395, 170–176 (2007).

82 Jose, A. M. & Hunter, C. P. Transport of sequence-specific RNA interference information between cells. Annu Rev Genet 41, 305–330, doi:10.1146/annurev.genet.41.110306.130216 (2007).

83 Egge, N. et al. Age-Onset Phosphorylation of a Minor Actin Variant Promotes Intestinal Barrier Dysfunction. Dev Cell 51, 587–601 e587, doi:10.1016/j.devcel.2019.11.001 (2019).

84 Kamath, R. S. & Ahringer, J. Genome-wide RNAi screening in Caenorhabditis elegans. Methods 30, 313–321, doi:10.1016/s1046-2023(03)00050-1 (2003).

85 Rual, J. F. et al. Toward improving Caenorhabditis elegans phenome mapping with an ORFeome-based RNAi library. Genome Res 14, 2162–2168, doi:10.1101/gr.2505604 (2004).

86 Brand, A. H. & Perrimon, N. Targeted gene expression as a means of altering cell fates and generating dominant phenotypes. development 118, 401–415 (1993).

87 Lee, T. & Luo, L. Mosaic analysis with a repressible cell marker for studies of gene function in neuronal morphogenesis. Neuron 22, 451–461 (1999).

88 Robinow, S. & White, K. Characterization and spatial distribution of the ELAV protein during Drosophila melanogaster development. J Neurobiol 22, 443–461, doi:10.1002/neu.480220503 (1991).

89 Han, S. K. et al. OASIS 2: online application for survival analysis 2 with features for the analysis of maximal lifespan and healthspan in aging research. Oncotarget 7, 56147 (2016).

90 Piper, M. D. & Partridge, L. Protocols to Study Aging in Drosophila. Methods Mol Biol 1478, 291–302, doi:10.1007/978-1-4939-6371-3_18 (2016).

91 Martins, R. R., McCracken, A. W., Simons, M. J., Henriques, C. M. & Rera, M. How to Catch a Smurf?– Ageing and Beyond… In vivo Assessment of Intestinal Permeability in Multiple Model Organisms. Bio-protocol 8 (2018).

92 Thomas, J.H. Genetic analysis of defecation in Caenorhabditis elegans. Genetics 124, 855–872 (1990).

