## Extended Data figures 1-6, Supplementary Tables 1-6 and legends for "Conserved neuropeptidergic regulation of intestinal integrity in invertebrate models of aging"

File includes Extended data figures 1-6, Supplementary Tables 1-6 and corresponding legends.

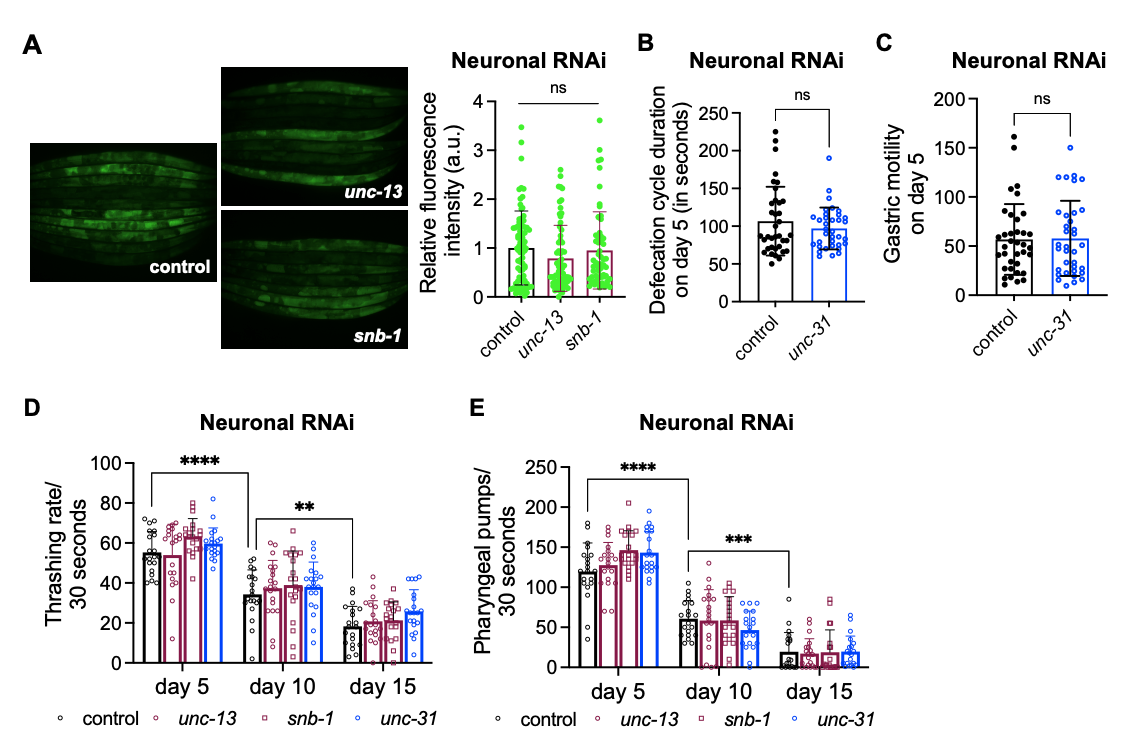
**Extended Data Fig. 1.** **Blocking neurotransmitter or neuropeptides release has no effects on gastric motility and muscle function.**

**(A)** Neuron-specific knockdown of *unc-13/Munc-13* or *snb-1/synaptobrevin* in neurons resulted in fluorescein intensities comparable to control animals. Represented data was quantified from n=1 out of 4 trails, N=77 (control), N=66 (*unc-13/Munc-13)*, N=60 (*snb-1/ synaptobrevin)*. ns *P>0.05, one-way ANOVA* with Dunnett's multiple comparisons test. See **Supplementary Table 3** for data of independent trials.

**(B)** Defecation frequency and **(C)** gastric motility of day 5 animals after neuron-specific RNAi of *unc-31/CAPS* initiated from day 1 of adulthood. Error bars are std. dev. of n=3, N>35. ns *P>0.05*, two-tailed t-test.

**(D)** Thrashing rate and **(E)** pharyngeal pumping after neuron-specific RNAi of *unc-13/Munc-13*, *snb-1/synaptobrevin* or *unc-31/CAPS*. Error bars are std. dev of n=2, N=20. **** *P<0.0001,* *** *P<0.001,* ** *P<0.01, two-way ANOVA* with Tukey's multiple comparisons test.

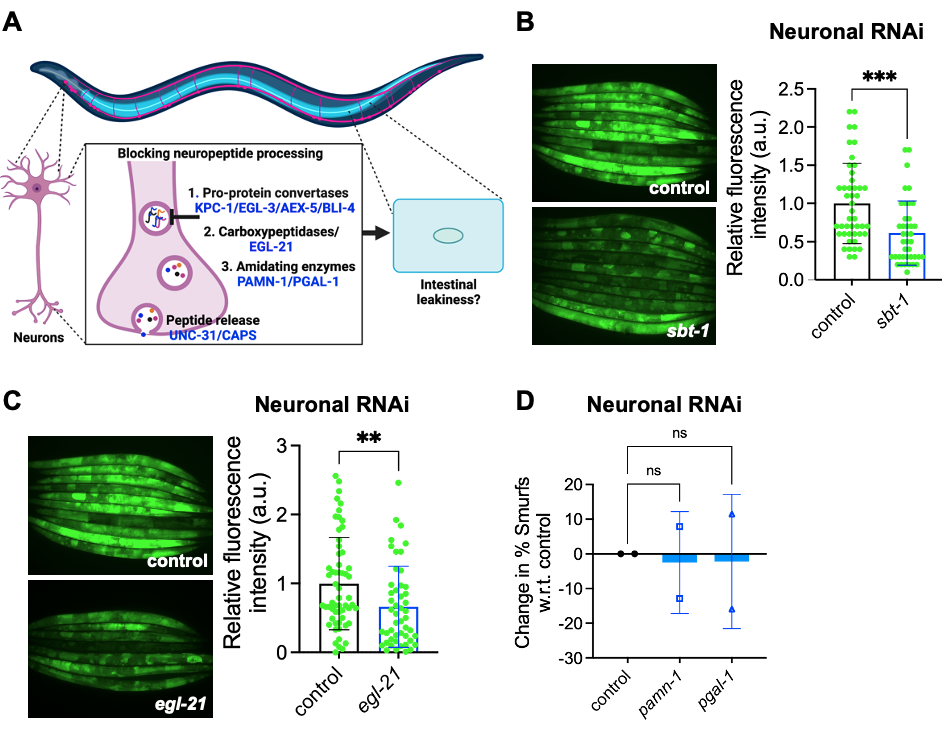

Extended Data Fig. 2. Neuropeptide processing modulates aging-evoked changes in intestinal leakiness

**(A)** Schematic of experimental approach for down regulating conserved sequential steps in peptide-processing to test their effect on intestinal leakiness in 15-day old nematodes.

**(B)** FluoroSmurf phenotype of EGL-3/PC2 Chaperone: *sbt-1*. Represented data quantified from n=1, N=45 (control), N=40 *(sbt-1/7B2),* out of 3 trails. *** *P<0.001*, two-tailed t-test.

**(C)** FluoroSmurf phenotype of carboxypeptidase E/*egl-21.* Represented data quantified from n=2, N=58 (control), N=49 (*egl-21/* carboxypeptidase E), out of 7 trails. ** *P<0.01*, two-tailed t-test.

For (B-C), also see **Supplementary Table 3** for data of independent trials.

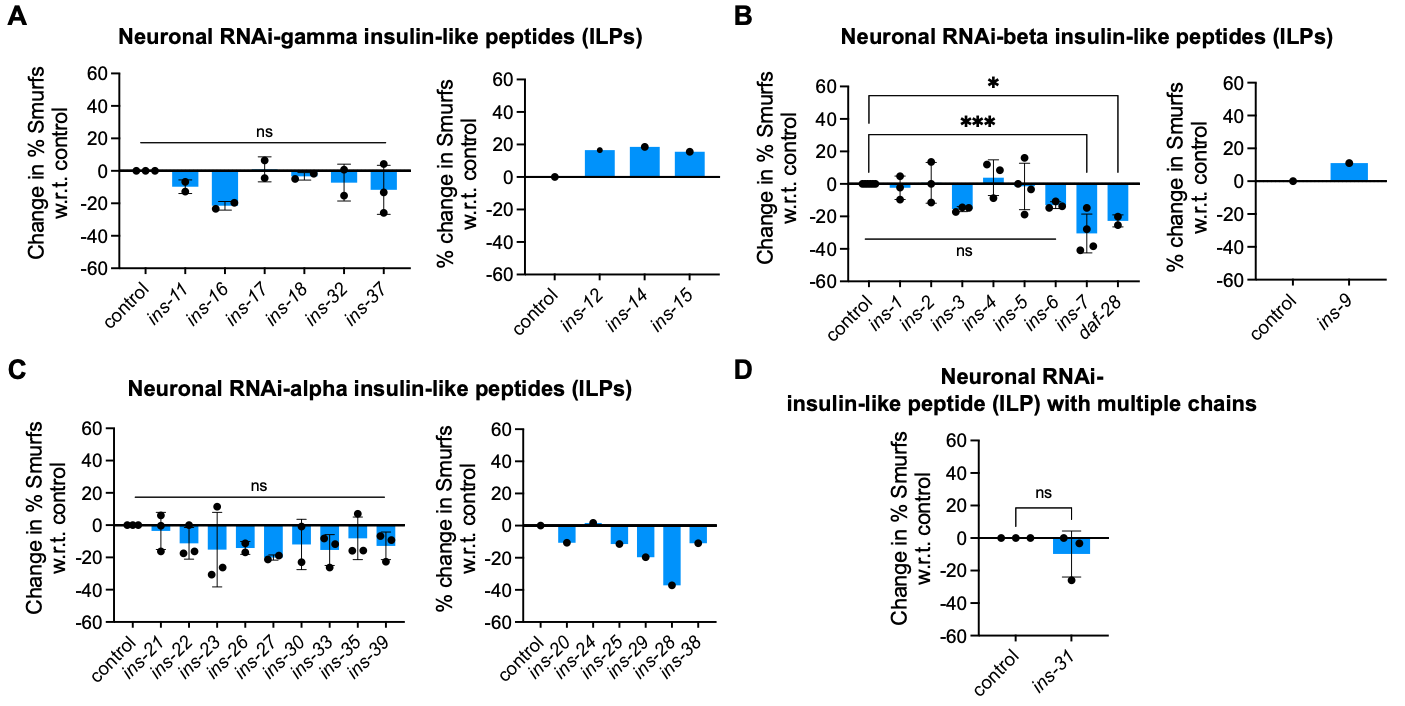
**Extended Data Fig. 3.** **Intestine-modifying roles of neuronal insulin-like peptides in *C. elegans*.**

Neuron-specific RNAi screen for 35 (out of 40) insulin-like peptide genes to test their effect on intestinal leakiness in 15-day old *C. elegans*. Insulin-like peptides genes have been classified based on their disulfide bonds [18, 19]. RNAi knockdown of individual insulin-like peptide gene was initiated on day 1 of adulthood. Intestinal leakiness represented as change in % Smurf w.r.t control (empty L4440 vector).

**(A)** Left panel- gamma *ins* gene tested more than two times; *ins-16* RNAi exhibited a trend for consistent decrease in %Smurf. Right panel- gamma *ins* gene knockdown showed a trend for increase in %Smurf, n=1.

**(B)** Left panel- beta *ins* genes tested more than two times; *ins-3* and *ins-6* RNAi exhibited a trend for consistent decrease in %Smurf, *ins-7* and *daf-28* RNAi exhibited a significant decrease in %Smurf. Right panel- beta *ins-9* RNAi showed a trend for increase in %Smurf, n=1.

**(C)** Left panel- alpha *ins* genes tested more than two times; *ins-27* and *ins-33* RNAi exhibited a trend for consistent decrease in %Smurf. Right panel- multiple alpha *ins* RNAi showed a trend for decrease in %Smurf, n=1.

*** *P<0.001*, * *P<0.05,* ns *P>0.05, one-way ANOVA* with Dunnett's multiple comparisons test.

**(D)** Neuron-specific RNAi of *ins-31,* the ILP gene with multiple insulin-like domains did not impact the age-related intestinal leakiness. n=3, ns *P>0.05*, two-tailed t-test.

See **Supplementary Table 4** for data of independent trials.

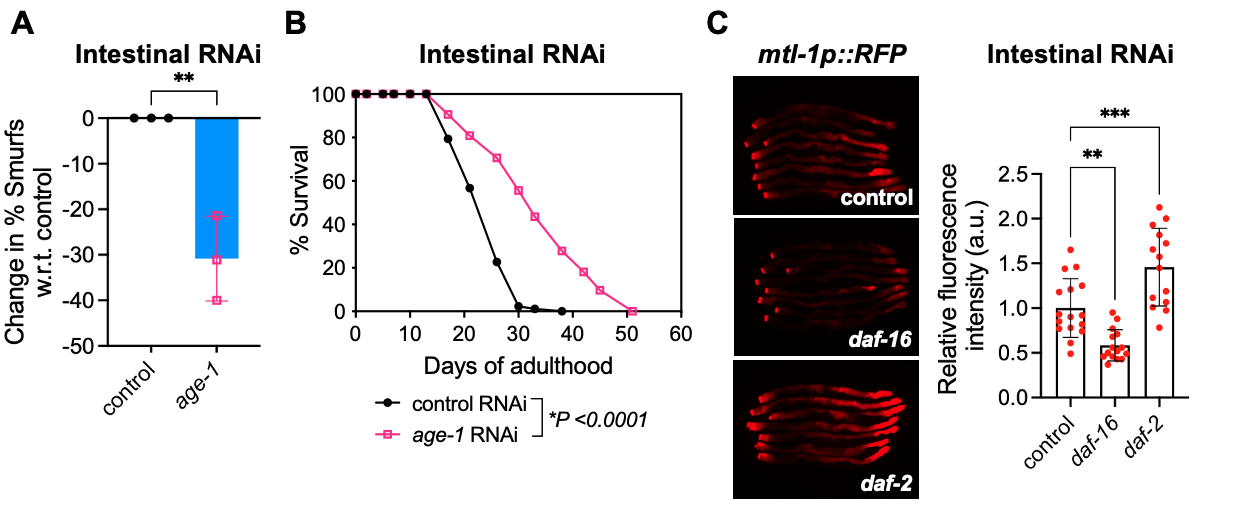

**Extended Data Fig. 4. Intestinal insulin signaling affects intestinal barrier integrity**

**(A)** HypoSmurf phenotype of intestine-specific KD of *age-1/PI3K*. Error bars are std. dev. for n=3 ** *P<0.001*, two-tailed t-test. See **Supplementary Table 1** for data of independent trials.

**(B)** Lifespan extension during intestine-specific knockdown of *age-1/PI3K (** *P< 0.0001,* log-rank test) initiated from day 1 of adulthood.

**(C)** MTL-1 expression during intestine-specific RNAi of *daf-1/InR or daf-16/FOXO.* RNAi was initiated on day 1 and imaged on day 5. Each dot represents relative fluorescent intensity in arbitrary units (a.u.) quantified from entire intestine region below the pharynx. Error bars are std. dev. of N>14, *** *P<0.001*, ** *P<0.01, one-way ANOVA* with Dunnett's multiple comparisons test. Data is representative of two independent trials.

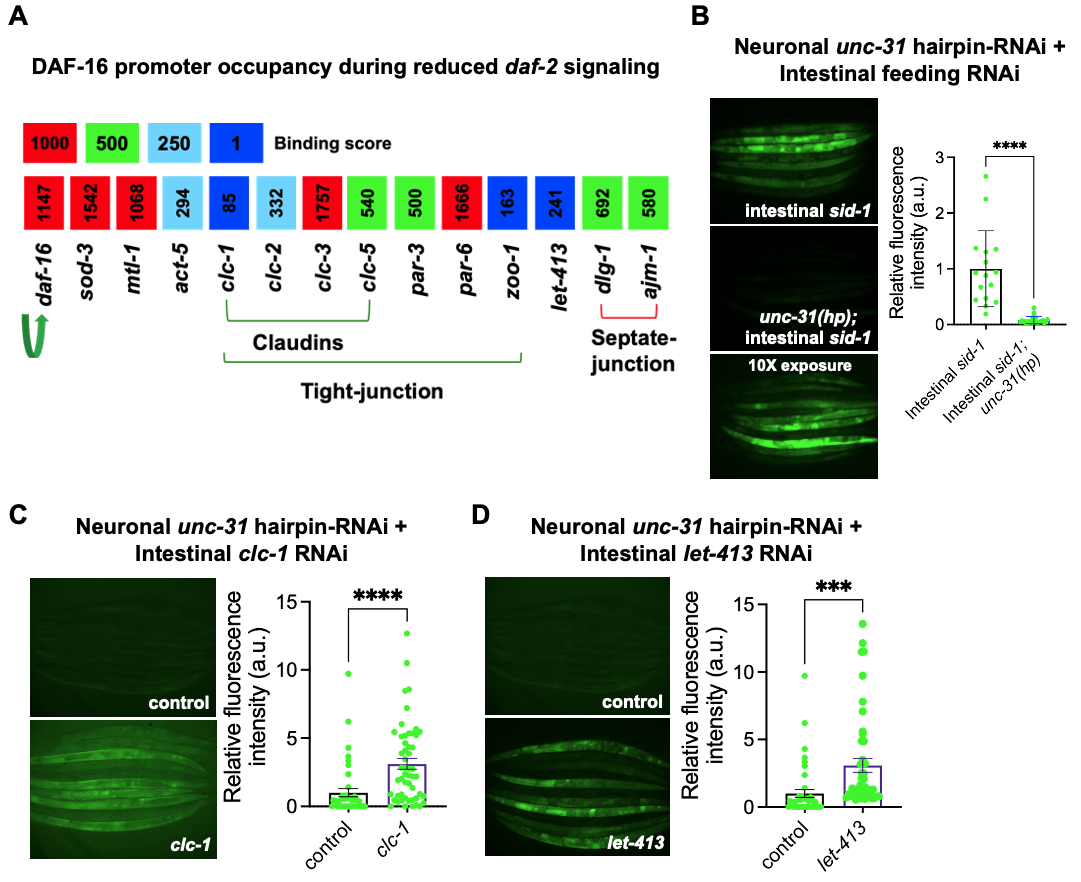

**Extended Data Fig. 5. Direct DAF-16/FOXO targets associated with epithelial barriers maintain intestinal integrity downstream of neuropeptide signals.**

**(A)** ChIP-Atlas data showing list of genes with enriched DAF-16/FOXO recruitment, binding peaks distributed within ±1kb with respect to the transcriptional start site (TSS). DAF-16 is known to bind its own promoter, creating a positive feedback loop (green arrow), and its bonafide targets, *mtl-1* and *sod-3*.

**(B)** HypoSmurf phenotype of transgenic strain generated by combining neuronal *unc-31/CAPS* hairpin RNAi with intestine-specific feeding RNAi transgenics exhibited HypoSmurf phenotype (lower panel image was acquired at 10X exposure time), similar to neuron-specific RNAi of *unc-31/CAPS.* Represented data quantified from n=1*,* N=16 (control)*,* N=24 *(act-5)*, out of 4 trails. **** *P<0.0001*, two-tailed t-test.

**(C)** Intestine-specific knockdown of *clc-1* (tight junction component) suppressed the HypoSmurf phenotype of neuronal *unc-31/CAPS* hairpin-RNAi transgenics. Represented data quantified from n=1*,* N=43 (control)*,* N=52 *(clc-1)*, out of 4 trails. **** *P<0.0001*, two-tailed t-test.

**(D)** Intestine-specific knockdown of *let-413* (a basolateral component) suppressed the HypoSmurf phenotype of neuronal *unc-31/CAPS* hairpin-RNAi transgenics. Represented data quantified from n=1*,* N=43 (control)*,* N=47 *(let-413)*, out of 2 trails. *** *P<0.001*, two-tailed t-test.

For (B-D), also see **Supplementary Table 1** for data of independent trials.

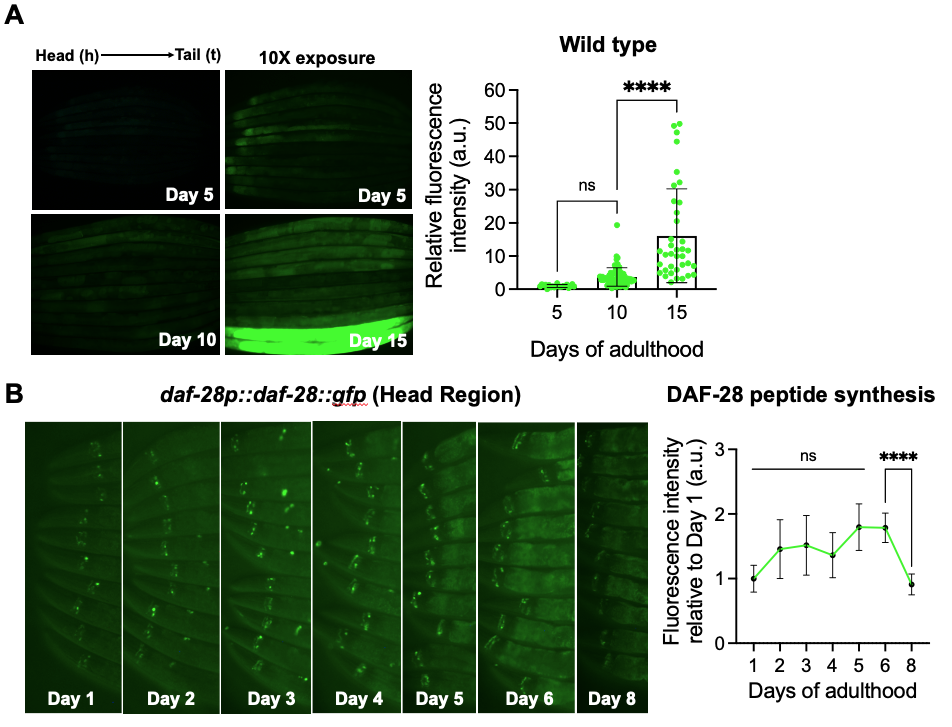

**Extended Data Fig. 6.** **Predominantly post-reproductive age-related increase in intestinal leakiness.**

**(A)** Modified FluoroSmurf method was implemented to quantify the extent of age-related leakiness. Wildtype (N2-CGC) animals were fed both blue and Fluorescein dye at indicated age-points and imaged for fluorescent dye accumulation at constant exposure time (5ms). Represented data was quantified from n=1, N=21 (day 5), N=72 (day 10), and N=37 (day 15), out of 3 trails. Each dot represents relative fluorescent intensity in arbitrary units (a.u.) quantified from entire body of each animal. Error bars are std. dev. **** *P<0.0001*, ns *P>0.05, one-way ANOVA* with Tukey's multiple comparisons test.

**(B)** The age-related synthesis of DAF-28 peptide measured as fluorescence intensity of GFP signals in the head neurons. Error bars are std. dev. of N>10. ***** P <0.0001,* ns *P>0.05, one-way ANOVA* with Tukey's multiple comparisons test. Data is representative of three independent trials.

**Supplementary Table 1: Details of Smurf and FluoroSmurf experiments**

| **Repeats for Smurf assays- Related to Figure 1-4. FUDR+ RNAi was initiated on Day 1 of adulthood** | | | | | | | | | | | | | | | | | | |
| --- | --- | --- | --- | --- | --- | --- | --- | --- | --- | --- | --- | --- | --- | --- | --- | --- | --- | --- |
| **Figure Reference** | **Strain identifier** | **Target tissue** | | | | **RNAi bacteria** | | **# of Non-Smurfs** | | | **# of Smurfs** | | | **Total # of animals** | | **% Smurf** | | **Change in % Smurf** |
| ***Related to Figure 1B*** | | | | | | | | | | | | | | | | | | |
| Repeat 1 | MAH677 | Neuron-specific  RNAi | | | | **control** | | **8** | | | **14** | | | **22** | | **63.6** | |  |
|  |  |  |  |  |  | *unc-31* | | 17 | | | 9 | | | 26 | | 34.6 | | -29.0 |
| Repeat 2 |  |  |  |  |  | **control** | | **3** | | | **35** | | | **38** | | **92.1** | |  |
|  |  |  |  |  |  | *unc-31* | | 12 | | | 29 | | | 41 | | 70.7 | | -21.4 |
| Repeat 3 |  |  |  |  |  | **control** | | **5** | | | **12** | | | **17** | | **70.6** | |  |
|  |  |  |  |  |  | *unc-31* | | 15 | | | 6 | | | 21 | | 28.6 | | -42.0 |
| Repeat 4 |  |  |  |  |  | **control** | | **6** | | | **14** | | | **20** | | **70.0** | |  |
|  |  |  |  |  |  | *unc-31* | | 18 | | | 13 | | | 31 | | 41.9 | | -28.1 |
| Repeat 5 |  |  |  |  |  | **control** | | **17** | | | **34** | | | **51** | | **66.7** | |  |
|  |  |  |  |  |  | *unc-31* | | 33 | | | 30 | | | 63 | | 47.6 | | -19.0 |
| Repeat 6 |  |  |  |  |  | **control** | | **20** | | | **56** | | | **76** | | **73.7** | |  |
|  |  |  |  |  |  | *unc-31* | | 38 | | | 29 | | | 67 | | 43.3 | | -30.4 |
| **Figure 1B**  **Quantified** Repeat 7 |  |  |  |  |  | **control** | | **4** | | | **28** | | | **32** | | **87.5** | |  |
|  |  |  |  |  |  | *unc-31* | | 16 | | | 17 | | | 33 | | 51.5 | | -36.0 |
| **Figure 1B**  **Quantified** Repeat 8 |  |  |  |  |  | **control** | | **13** | | | **28** | | | **41** | | **68.3** | |  |
|  |  |  |  |  |  | *unc-31* | | 16 | | | 21 | | | 37 | | 56.8 | | -11.5 |
| ***Related to Figure 1C*** | | | | | | | | | | | | | | | | | | |
| Repeat 1 | MAH677 | Neuron-specific  RNAi | | | | **control** | | **8** | | | **14** | | | **22** | | **63.6** | |  |
|  |  |  |  |  |  | *kpc-1* | | 12 | | | 7 | | | 19 | | 36.8 | | -26.8 |
|  |  |  |  |  |  | *egl-3* | | 33 | | | 3 | | | 36 | | 8.3 | | -55.3 |
|  |  |  |  |  |  | *aex-5* | | 11 | | | 28 | | | 39 | | 71.8 | | + 8.2 |
|  |  |  |  |  |  | *bli-4* | | 9 | | | 17 | | | 26 | | 65.4 | | + 1.7 |
| Repeat 2 |  |  |  |  |  | **control** | | **3** | | | **35** | | | **38** | | **92.1** | |  |
|  |  |  |  |  |  | *kpc-1* | | 12 | | | 17 | | | 29 | | 58.6 | | -33.5 |
|  |  |  |  |  |  | *egl-3* | | 18 | | | 17 | | | 35 | | 48.6 | | -43.5 |
|  |  |  |  |  |  | *aex-5* | | 5 | | | 44 | | | 49 | | 89.8 | | - 2.3 |
|  |  |  |  |  |  | *bli-4* | | 9 | | | 23 | | | 32 | | 71.9 | | -20.2 |
| Repeat 3 |  |  |  |  |  | **control** | | **11** | | | **18** | | | **29** | | **62.1** | |  |
|  |  |  |  |  |  | *kpc-1* | | 8 | | | 21 | | | 29 | | 72.4 | | +10.3 |
|  |  |  |  |  |  | *egl-3* | | 13 | | | 19 | | | 32 | | 59.4 | | - 2.7 |
|  |  |  |  |  |  | *aex-5* | | 7 | | | 18 | | | 25 | | 72.0 | | + 9.9 |
|  |  |  |  |  |  | *bli-4* | | 11 | | | 19 | | | 30 | | 63.3 | | + 1.3 |
| Repeat 4 |  |  |  |  |  | **control** | | **4** | | | **28** | | | **32** | | **87.5** | |  |
|  |  |  |  |  |  | *egl-3* | | 46 | | | 10 | | | 56 | | 17.9 | | -69.6 |
|  |  |  |  |  |  | *aex-5* | | 27 | | | 25 | | | 52 | | 48.1 | | -39.4 |
| Repeat 5 |  |  |  |  |  | **control** | | **29** | | | **37** | | | **66** | | **56.1** | |  |
|  |  |  |  |  |  | *egl-3* | | 44 | | | 11 | | | 55 | | 20.0 | | -36.1 |
|  |  |  |  |  |  | *aex-5* | | 22 | | | 19 | | | 41 | | 46.3 | | - 9.7 |
| Repeat 6 |  |  |  |  |  | **control** | | **17** | | | **34** | | | **51** | | **66.7** | |  |
|  |  |  |  |  |  | *kpc-1* | | 47 | | | 31 | | | 78 | | 39.7 | | -26.9 |
|  |  |  |  |  |  | *egl-3* | | 48 | | | 19 | | | 67 | | 28.4 | | -38.3 |
|  |  |  |  |  |  | *aex-5* | | 43 | | | 20 | | | 63 | | 31.7 | | -34.9 |
|  |  |  |  |  |  | *bli-4* | | 13 | | | 40 | | | 53 | | 75.5 | | + 8.8 |
| Repeat 7  **Quantified**  **Figure 1C** |  |  |  |  |  | **control** | | **15** | | | **32** | | | **47** | | **68.1** | |  |
|  |  |  |  |  |  | *kpc-1* | | 13 | | | 15 | | | 28 | | 53.6 | | -14.5 |
|  |  |  |  |  |  | *egl-3* | | 26 | | | 10 | | | 36 | | 27.8 | | -40.3 |
|  |  |  |  |  |  | *aex-5* | | 10 | | | 18 | | | 28 | | 64.3 | | - 3.8 |
|  |  |  |  |  |  | *bli-4* | | 6 | | | 23 | | | 29 | | 79.3 | | +11.2 |
| ***Related to Figure 1D*** | | | | | | | | | | | | | | | | | | |
| Repeat 1 | Wild type | |  | | | **control** | | **22** | | | **40** | | | **62** | | **64.5** | |  |
|  | MAH1094 | | Neuronal *egl-3 OE* | | |  |  | 2 | | | 48 | | | 50 | | 96.0 | | +31.5 |
| Repeat 2 | Wild type | |  | | |  |  | **18** | | | **29** | | | **47** | | **61.7** | |  |
|  | MAH1094 | | Neuronal *egl-3 OE* | | |  |  | 4 | | | 42 | | | 46 | | 91.3 | | +29.6 |
| Repeat 3 | Wild type | |  | | |  |  | **21** | | | **46** | | | **67** | | **68.7** | |  |
|  | MAH1094 | | Neuronal *egl-3 OE* | | |  |  | 7 | | | 48 | | | 55 | | 87.3 | | +18.6 |
| Repeat 4 | Wild type | |  | | |  |  | **19** | | | **20** | | | **39** | | **51.3** | |  |
|  | MAH1094 | | Neuronal *egl-3 OE* | | |  |  | 18 | | | 49 | | | 67 | | 73.1 | | +21.9 |
| Repeat 5  **Figure 1D** **Quantified** | Wild type | |  | | |  |  | **18** | | | **39** | | | **57** | | **68.4** | |  |
|  | MAH1094 | | Neuronal *egl-3 OE* | | |  |  | 11 | | | 47 | | | 58 | | 81.0 | | +12.6 |
| ***Related to Figure S1A*** | | | | | | | | | | | | | | | | | | |
| Repeat 1 | MAH677 | | Neuron-specific  RNAi | | | **control** | | **7** | | | **21** | | | **28** | | **75.0** | |  |
|  |  |  |  |  |  | *unc-13* | | 5 | | | 35 | | | 40 | | 87.5 | | +12.5 |
|  |  |  |  |  |  | *snb-1* | | 5 | | | 13 | | | 18 | | 72.2 | | - 2.8 |
| Repeat 2 |  |  |  |  |  | **control** | | **13** | | | **28** | | | **41** | | **68.3** | |  |
|  |  |  |  |  |  | *unc-13* | | 18 | | | 28 | | | 46 | | 60.9 | | - 7.4 |
|  |  |  |  |  |  | *snb-1* | | 24 | | | 26 | | | 50 | | 52.0 | | -16.3 |
| Repeat 3 |  |  |  |  |  | **control** | | **23** | | | **21** | | | **44** | | **47.7** | |  |
|  |  |  |  |  |  | *unc-13* | | 27 | | | 25 | | | 52 | | 48.1 | | + 0.3 |
|  |  |  |  |  |  | *snb-1* | | 30 | | | 17 | | | 47 | | 36.2 | | -11.6 |
| **Figure S1A**  **Quantified**  Repeat 4 |  |  |  |  |  | **control** | | **20** | | | **56** | | | **76** | | **73.7** | |  |
|  |  |  |  |  |  | *unc-13* | | 35 | | | 33 | | | 68 | | 48.5 | | -25.2 |
|  |  |  |  |  |  | *snb-1* | | 32 | | | 37 | | | 69 | | 53.6 | | -20.1 |
| ***Related to Figure S2B*** | | | | | | | | | | | | | | | | | | |
| Repeat 1 | MAH677 | | Neuron-specific  RNAi | | | **control** | | **17** | | | **34** | | | **51** | | **66.7** | |  |
|  |  |  |  |  |  | *sbt-1* | | 33 | | | 18 | | | 51 | | 35.3 | | -31.4 |
| Repeat 2 |  |  |  |  |  | **control** | | **20** | | | **56** | | | **76** | | **73.7** | |  |
|  |  |  |  |  |  | *sbt-1* | | 38 | | | 27 | | | 65 | | 41.5 | | -32.1 |
| Repeat 3  **Figure S2B** Quantified |  |  |  |  |  | **control** | | **15** | | | **32** | | | **47** | | **68.1** | |  |
|  |  |  |  |  |  | *sbt-1* | | 22 | | | 18 | | | 40 | | 45.0 | | -23.1 |
| ***Related to Figure S2C*** | | | | | | | | | | | | | | | | | | |
| Repeat 1 | MAH677 | | Neuron-specific  RNAi | | | **control** | | **3** | | | **12** | | | **15** | | **80.0** | |  |
|  |  |  |  |  |  | *egl-21* | | 13 | | | 10 | | | 23 | | 43.5 | | -36.5 |
| Repeat 2 |  |  |  |  |  | **control** | | **13** | | | **25** | | | **38** | | **65.8** | |  |
|  |  |  |  |  |  | *egl-21* | | 22 | | | 23 | | | 45 | | 51.1 | | -14.7 |
| Repeat 3 |  |  |  |  |  | **control** | | **10** | | | **10** | | | **20** | | **50.0** | |  |
|  |  |  |  |  |  | *egl-21* | | 31 | | | 11 | | | 42 | | 26.2 | | -23.1 |
| Repeat 4 |  |  |  |  |  | **control** | | **6** | | | **14** | | | **20** | | **70.0** | |  |
|  |  |  |  |  |  | *egl-21* | | 25 | | | 11 | | | 36 | | 30.6 | | -39.4 |
| Repeat 5 |  |  |  |  |  | **control** | | **17** | | | **34** | | | **51** | | **66.7** | |  |
|  |  |  |  |  |  | *egl-21* | | 45 | | | 22 | | | 67 | | 32.8 | | -33.8 |
| Repeat 6  **Figure S2C** Quantified |  |  |  |  |  | **control** | | **4** | | | **28** | | | **32** | | **87.5** | |  |
|  |  |  |  |  |  | *egl-21* | | 26 | | | 8 | | | 34 | | 23.5 | | -64.0 |
| Repeat 7  **Figure S2C** Quantified |  |  |  |  |  | **control** | | **15** | | | **32** | | | **47** | | **68.1** | |  |
|  |  |  |  |  |  | *egl-21* | | 15 | | | 15 | | | 30 | | 50.0 | | -18.1 |
| ***Related to Figure S2D*** | | | | | | | | | | | | | | | | | | |
| Repeat 1 | MAH677 | | Neuron-specific  RNAi | | | **control** | | **7** | | | **21** | | | **28** | | **75.0** | |  |
|  |  |  |  |  |  | *pamn-1* | | 6 | | | 29 | | | 35 | | 82.9 | | + 7.9 |
|  |  |  |  |  |  | *pgal-1* | | 5 | | | 32 | | | 37 | | 86.5 | | +11.5 |
| Repeat 2 |  |  |  |  |  | **control** | | **3** | | | **30** | | | **33** | | **90.9** | |  |
|  |  |  |  |  |  | *pamn-1* | | 9 | | | 32 | | | 41 | | 78.0 | | -12.9 |
|  |  |  |  |  |  | *pgal-1* | | 9 | | | 27 | | | 36 | | 75.0 | | -15.9 |
| ***Related to Figure 2B*** | | | | | | | | | | | | | | | | | | |
| Repeat 1 | MAH677 | | Neuron-specific  RNAi | | | **control** | | **17** | | | **34** | | | **51** | | **66.7** | |  |
|  |  |  |  |  |  | *daf-28* | | 39 | | | 34 | | | 73 | | 46.6 | | -20.1 |
| Repeat 2 |  |  |  |  |  | **control** | | **17** | | | **36** | | | **53** | | **67.9** | |  |
|  |  |  |  |  |  | *daf-28* | | 28 | | | 31 | | | 59 | | 52.5 | | -15.4 |
| Repeat 3  **Figure 2B** Quantified |  |  |  |  |  | **control** | | **20** | | | **56** | | | **76** | | **73.7** | |  |
|  |  |  |  |  |  | *daf-28* | | 31 | | | 29 | | | 60 | | 48.3 | | -25.4 |
| ***Related to Figure 2C*** | | | | | | | | | | | | | | | | | | |
| Repeat 1 | MAH677 | | Neuron-specific  RNAi | | | **control** | | **17** | | | **42** | | | **59** | | **71.2** | |  |
|  |  |  |  |  |  | *ins-7* | | 39 | | | 19 | | | 58 | | 32.8 | | -38.4 |
| Repeat 2 |  |  |  |  |  | **control** | | **4** | | | **24** | | | **28** | | **85.7** | |  |
|  |  |  |  |  |  | *ins-7* | | 9 | | | 22 | | | 31 | | 71.0 | | -14.7 |
| Repeat 3 |  |  |  |  |  | **control** | | **13** | | | **25** | | | **38** | | **65.8** | |  |
|  |  |  |  |  |  | *ins-7* | | 18 | | | 11 | | | 29 | | 37.9 | | -27.9 |
| Repeat 4 |  |  |  |  |  | **control** | | **10** | | | **10** | | | **20** | | **50.0** | |  |
|  |  |  |  |  |  | *ins-7* | | 19 | | | 8 | | | 27 | | 29.6 | | -20.4 |
| Repeat 5 |  |  |  |  |  | **control** | | **0** | | | **16** | | | **16** | | **100.0** | |  |
|  |  |  |  |  |  | *ins-7* | | 12 | | | 21 | | | 33 | | 63.6 | | -36.4 |
| Repeat 6 |  |  |  |  |  | **control** | | **6** | | | **14** | | | **20** | | **70.0** | |  |
|  |  |  |  |  |  | *ins-7* | | 48 | | | 21 | | | 69 | | 30.4 | | -39.6 |
| Repeat 7 |  |  |  |  |  | **control** | | **19** | | | **21** | | | **40** | | **52.5** | |  |
|  |  |  |  |  |  | *ins-7* | | 41 | | | 20 | | | 61 | | 32.8 | | -19.7 |
| Repeat 8 |  |  |  |  |  | **control** | | **4** | | | **53** | | | **57** | | **93.0** | |  |
|  |  |  |  |  |  | *ins-7* | | 21 | | | 35 | | | 56 | | 62.5 | | -30.5 |
| Repeat 9  **Figure 2C** Quantified |  |  |  |  |  | **control** | | **17** | | | **36** | | | **53** | | **67.9** | |  |
|  |  |  |  |  |  | *ins-7* | | 28 | | | 32 | | | 60 | | 53.3 | | -14.6 |
| ***Related to Figure 2D*** | | | | | | | | | | | | | | | | | | |
| Repeat 1 | MAH1256 | | Neuronal *empty vector* | | | **control** | | **24** | | | **17** | | | **41** | | **41.5** | | **+18.5** |
| Repeat 2 | MAH1257 | |  |  |  |  |  | **30** | | | **15** | | | **45** | | **33.3** | | **+10.5** |
| Repeat 3 | MAH1258 | |  |  |  |  |  | **37** | | | **11** | | | **48** | | **22.9** | |  |
| Repeat 4 | MAH1259 | |  |  |  |  |  | **30** | | | **18** | | | **48** | | **37.5** | | **+14.6** |
| Repeat 1 | MAH1264 | | Neuronal *daf-28 OE* | | |  |  | 19 | | | 20 | | | 39 | | 51.3 | | +28.4 |
| Repeat 2 | MAH1265 | |  |  |  |  |  | 19 | | | 19 | | | 38 | | 50.0 | | +27.1 |
| Repeat 5 | MAH1257 | | Neuronal *empty vector* | | |  |  | **19** | | | **11** | | | **30** | | **36.7** | | **+10.4** |
| Repeat 6 **Figure 2D**  Quantified | MAH1258 | |  |  |  |  |  | **28** | | | **10** | | | **38** | | **26.3** | |  |
| Repeat 7 **Figure 2D** Quantified | MAH1259 | |  |  |  |  |  | **14** | | | **10** | | | **24** | | **41.7** | | **+15.4** |
| Repeat 3 | MAH1265 | | Neuronal *daf-28 OE* | | |  |  | 20 | | | 22 | | | 42 | | 52.4 | | +26.1 |
| Repeat 4 **Figure 2D** Quantified | MAH1264 | |  |  |  |  |  | 7 | | | 16 | | | 23 | | 69.6 | | +43.2 |
| Repeat 5 **Figure 2D** Quantified | MAH1264 | |  |  |  |  |  | 7 | | | 21 | | | 28 | | 75.0 | | +48.7 |
| Repeat 6 **Figure 2D** Quantified | MAH1265 | |  |  |  |  |  | 9 | | | 15 | | | 24 | | 62.5 | | +36.2 |
| ***Related to Figure 2E*** | | | | | | | | | | | | | | | | | | |
| Repeat 1 | MAH1257 | | | Neuronal *empty vector* | | **control** | | **30** | | | **15** | | | **45** | | **33.3** | | **+10.4** |
| Repeat 2 | MAH1258 | | |  |  |  |  | **37** | | | **11** | | | **48** | | **22.9** | |  |
| Repeat 3 **Figure 2E** Quantified | MAH1256 | | |  |  |  |  | **24** | | | **17** | | | **41** | | **41.5** | | **+18.5** |
| Repeat 4 **Figure 2E** Quantified | MAH1259 | | |  |  |  |  | **30** | | | **18** | | | **48** | | **37.5** | | **+14.6** |
| Repeat 1 | MAH1262 | | | Neuronal *ins-7 OE* | |  |  | 27 | | | 20 | | | 47 | | 42.6 | | +19.7 |
| Repeat 2 **Figure 2E** Quantified | MAH1260 | | |  |  |  |  | 21 | | | 25 | | | 46 | | 54.3 | | +31.5 |
| Repeat 3 **Figure 2E** Quantified | MAH1261 | | |  |  |  |  | 23 | | | 26 | | | 49 | | 53.1 | | +30.1 |
| ***Related to Figure 3B*** | | | | | | | | | | | | | | | | | | |
| Repeat 1 | MAH728 | | | Intestine-specific  RNAi | | | **control** | **0** | | | **12** | | | **12** | | **100.0** | |  |
|  |  |  |  |  |  |  | *daf-2* | 4 | | | 4 | | | 8 | | 50.0 | | -50.0 |
| Repeat 2 |  |  |  |  |  |  | **control** | **3** | | | **14** | | | **17** | | **82.4** | |  |
|  |  |  |  |  |  |  | *daf-2* | 16 | | | 12 | | | 28 | | 42.9 | | -39.5 |
| Repeat 3 |  |  |  |  |  |  | **control** | **7** | | | **30** | | | **37** | | **81.1** | |  |
|  |  |  |  |  |  |  | *daf-2* | 18 | | | 11 | | | 29 | | 37.9 | | -43.2 |
| Repeat 4 |  |  |  |  |  |  | **control** | **4** | | | **40** | | | **44** | | **90.9** | |  |
|  |  |  |  |  |  |  | *daf-2* | 13 | | | 31 | | | 44 | | 70.5 | | -20.4 |
| Repeat 5 |  |  |  |  |  |  | **control** | **11** | | | **21** | | | **32** | | **65.6** | |  |
|  |  |  |  |  |  |  | *daf-2* | 29 | | | 4 | | | 33 | | 12.1 | | -53.5 |
| Repeat 6 |  |  |  |  |  |  | **control** | **0** | | | **19** | | | **19** | | **100.0** | |  |
|  |  |  |  |  |  |  | *daf-2* | 19 | | | 24 | | | 43 | | 55.8 | | -44.2 |
| Repeat 7 |  |  |  |  |  |  | **control** | **2** | | | **16** | | | **18** | | **88.9** | |  |
|  |  |  |  |  |  |  | *daf-2* | 30 | | | 5 | | | 35 | | 14.3 | | -74.6 |
| Repeat 8 |  |  |  |  |  |  | **control** | **4** | | | **15** | | | **19** | | **78.9** | |  |
|  |  |  |  |  |  |  | *daf-2* | 22 | | | 20 | | | 42 | | 47.6 | | -31.3 |
| Repeat 9  **Figure 3B** **Quantified** |  |  |  |  |  |  | **control** | **14** | | | **17** | | | **31** | | **54.8** | |  |
|  |  |  |  |  |  |  | *daf-2* | 24 | | | 5 | | | 29 | | 17.2 | | -37.6 |
| ***Related to Figure S4A*** | | | | | | | | | | | | | | | | | | |
| Repeat 1 | MAH728 | | | Intestine-specific  RNAi | | | **control** | **0** | | | **12** | | | **12** | | **100.0** | |  |
|  |  |  |  |  |  |  | *age-1* | 3 | | | 11 | | | 14 | | 78.6 | | -21.4 |
| Repeat 2 |  |  |  |  |  |  | **control** | **3** | | | **14** | | | **17** | | **82.4** | |  |
|  |  |  |  |  |  |  | *age-1* | 15 | | | 11 | | | 26 | | 42.3 | | -40.1 |
| Repeat 3 |  |  |  |  |  |  | **control** | **7** | | | **30** | | | **37** | | **81.1** | |  |
|  |  |  |  |  |  |  | *age-1* | 13 | | | 13 | | | 26 | | 50.0 | | -31.1 |
| ***Related to Figure S5B*** | | | | | | | | | | | | | | | | | | |
| Repeat 1 | MAH728 | | | Intestine-specific  RNAi | | | control | **28** | | | **23** | | | **51** | | **45.1** | |  |
|  | MAH1093 | | | Neuron-specific  *unc-31* RNAi +  Intestine-specific  RNAi | | |  | 26 | | | 15 | | | 41 | | 36.6 | | -8.5 |
| Repeat 2 | MAH728 | | | Intestine-specific  RNAi | | | control | **18** | | | **18** | | | **36** | | **50.0** | |  |
|  | MAH1093 | | | Neuron-specific  *unc-31* RNAi +  Intestine-specific  RNAi | | |  | 28 | | | 11 | | | 39 | | 28.2 | | -21.8 |
| Repeat 3 | MAH728 | | | Intestine-specific  RNAi | | | control | **5** | | | **9** | | | **14** | | **64.3** | |  |
|  | MAH1093 | | | Neuron-specific  *unc-31* RNAi +  Intestine-specific  RNAi | | |  | 27 | | | 11 | | | 38 | | 28.9 | | -35.3 |
| Repeat 4  **Figure S6A** Quantified | MAH728 | | | Intestine-specific  RNAi | | | control | **4** | | | **12** | | | **16** | | **75.0** | |  |
|  | MAH1093 | | | Neuron-specific  *unc-31* RNAi +  Intestine-specific  RNAi | | |  | 12 | | | 11 | | | 23 | | 47.8 | | -27.2 |
| ***Related to Figure 3H*** | | | | | | | | | | | | | | | | | | |
| Repeat 1 | MAH1093 | | | Neuron-specific  *unc-31* RNAi +  Intestine-specific  RNAi | | | **control** | **35** | | | **11** | | | **46** | | **23.9** | |  |
|  |  |  |  |  |  |  | *daf-16* | 7 | | | 32 | | | 39 | | 82.1 | | +58.1 |
| Repeat 2 |  |  |  |  |  |  | **control** | **27** | | | **11** | | | **38** | | **28.9** | |  |
|  |  |  |  |  |  |  | *daf-16* | 27 | | | 24 | | | 51 | | 47.1 | | +18.1 |
| Repeat 3 |  |  |  |  |  |  | **control** | **26** | | | **15** | | | **41** | | **36.6** | |  |
|  |  |  |  |  |  |  | *daf-16* | 8 | | | 12 | | | 20 | | 60.0 | | +23.4 |
| Repeat 4 **Figure 3H** Quantified |  |  |  |  |  |  | **control** | **34** | | | **9** | | | **43** | | **20.9** | |  |
|  |  |  |  |  |  |  | *daf-16* | 14 | | | 29 | | | 43 | | 67.4 | | +46.5 |
| Repeat 5  **Figure 3H** Quantified |  |  |  |  |  |  | **control** | **12** | | | **11** | | | **23** | | **47.8** | |  |
|  |  |  |  |  |  |  | *daf-16* | 13 | | | 19 | | | 32 | | 59.4 | | +11.5 |
| ***Related to Figure 3I*** | | | | | | | | | | | | | | | | | | |
| Repeat 1 | MAH1093 | | | Neuron-specific  *unc-31* RNAi +  Intestine-specific  RNAi | | | **control** | **35** | | **11** | | | **46** | **23.9** | | | |  |
|  |  |  |  |  |  |  | *act-5* | 29 | | 30 | | | 59 | 50.8 | | | | +26.9 |
| Repeat 2  **Figure 3I**  **Quantified** |  |  |  |  |  |  | **control** | **34** | | **9** | | | **43** | **20.9** | | | |  |
|  |  |  |  |  |  |  | *act-5* | 24 | | 12 | | | 36 | 33.3 | | | | +12.4 |
| ***Related to Figure 3J*** | | | | | | | | | | | | | | | | | | |
| Repeat 1  **Figure 3J**  **Quantified** | MAH1093 | | | Neuron-specific  *unc-31* RNAi +  Intestine-specific  RNAi | | | **control** | **12** | | **11** | | | **23** | **47.8** | | | |  |
|  |  |  |  |  |  |  | *ajm-1* | 13 | | 15 | | | 28 | 53.6 | | | | +5.7 |
| ***Related to Figure S5C*** | | | | | | | | | | | | | | | | | | |
| Repeat 1 | MAH1093 | | | Neuron-specific  *unc-31* RNAi +  Intestine-specific  RNAi | | | **control** | **35** | | **11** | | | **46** | **23.9** | | | |  |
|  |  |  |  |  |  |  | *clc-1* | 29 | | 30 | | | 59 | 50.8 | | | | +26.9 |
| Repeat 2 |  |  |  |  |  |  | **control** | **28** | | **11** | | | **39** | **28.2** | | | |  |
|  |  |  |  |  |  |  | *clc-1* | 20 | | 15 | | | 35 | 42.9 | | | | +14.7 |
| Repeat 3 |  |  |  |  |  |  | **control** | **27** | | **11** | | | **38** | **28.9** | | | |  |
|  |  |  |  |  |  |  | *clc-1* | 37 | | 22 | | | 59 | 37.3 | | | | +8.3 |
| Repeat 4 **Figure S5C Quantified** |  |  |  |  |  |  | **control** | **34** | | **9** | | | **43** | **20.9** | | | |  |
|  |  |  |  |  |  |  | *clc-1* | 30 | | 22 | | | 52 | 42.3 | | | | +21.4 |
| ***Related to Figure S5D*** | | | | | | | | | | | | | | | | | | |
| Repeat 1 | MAH1093 | | | Neuron-specific  *unc-31* RNAi +  Intestine-specific  RNAi | | | **control** | **35** | | **11** | | | **46** | **23.9** | | | |  |
|  |  |  |  |  |  |  | *let-413* | 44 | | 23 | | | 67 | 34.3 | | | | +10.4 |
| Repeat 2  **Figure S5D Quantified** |  |  |  |  |  |  | **control** | **34** | | **9** | | | **43** | **20.9** | | | |  |
|  |  |  |  |  |  |  | *let-413* | 26 | | 21 | | | 47 | 44.7 | | | | +23.8 |
| ***Related to Figure 4F*** | | | | | | | | | | | | | | | | | | |
| Repeat 1 | Wild type | | |  | | | **control** | **11** | | **54** | | | **65** | **83.1** | | | |  |
|  | CF1903 | | | Germline-less | | |  | 49 | | 36 | | | 85 | 42.4 | | | | -40.7 |
| Repeat 2 | Wild type | | |  | | |  | **2** | | **50** | | | **52** | **96.2** | | | |  |
|  | CF1903 | | | Germline-less | | |  | 29 | | 30 | | | 59 | 50.8 | | | | -45.3 |
| Repeat 3 | Wild type | | |  | | |  | **20** | | **11** | | | **31** | **35.5** | | | |  |
|  | CF1903 | | | Germline-less | | |  | 34 | | 13 | | | 47 | 27.7 | | | | -7.8 |
| Repeat 4 | Wild type | | |  | | |  | **18** | | **20** | | | **38** | **52.6** | | | |  |
|  | CF1903 | | | Germline-less | | |  | 32 | | 20 | | | 52 | 38.5 | | | | -14.2 |
| Repeat 5  **Figure 4F Quantified** | Wild type | | |  | | |  | **17** | | **26** | | | **43** | **60.5** | | | |  |
|  | CF1903 | | | Germline-less | | |  | 59 | | 8 | | | 67 | 11.9 | | | | -48.5 |
| ***Related to Figure 4G*** | | | | | | | | | | | | | | | | | | |
| Repeat 1 | KUM11 | | | Germline-less | | | **control** | **10** | | **10** | | | **20** | **50.0** | | | |  |
|  | KUM12 | | | Germline-less + Neuronal *daf-28 OE* | | |  | 0 | | 10 | | | 10 | 100.0 | | | | +50.0 |
| Repeat 2 | KUM11 | | | Germline-less | | |  | **10** | | **6** | | | **16** | **37.5** | | | |  |
|  | KUM12 | | | Germline-less + Neuronal *daf-28 OE* | | |  | 19 | | 19 | | | 38 | 50.0 | | | | +12.5 |
| Repeat 3  **Figure 4G Quantified** | KUM11 | | | Germline-less | | |  | **8** | | **0** | | | **8** | **0.0** | | | |  |
|  | KUM12 | | | Germline-less + Neuronal *daf-28 OE* | | |  | 7 | | 10 | | | 17 | 58.8 | | | | +58.8 |
| **Figure Reference** | **Strain identifier** | | | **Age-point** | **RNAi bacteria** | | **Target tissue** | | **# of Non-Smurf** | | | **# of Smurf** | | | **Total # of animals** | | **% Smurf** | |
| ***Related to Figure S6As*** | | | | | | | | | | | | | | | | | | |
| Repeat 1 | Wild type | | | Day 5 | control | | Systemic | | **9** | | | **1** | | | **10** | | **10.0** | |
|  |  |  |  | Day 10 |  |  |  |  | 8 | | | 2 | | | 10 | | 20.0 | |
|  |  |  |  | Day 15 |  |  |  |  | 1 | | | 4 | | | 5 | | 80.0 | |
| Repeat 2 | Wild type | | | Day 5 |  |  |  |  | **58** | | | **4** | | | **62** | | **6.5** | |
|  |  |  |  | Day 10 |  |  |  |  | 15 | | | 7 | | | 22 | | 31.8 | |
|  |  |  |  | Day 15 |  |  |  |  | 3 | | | 5 | | | 8 | | 62.5 | |
| Repeat 3  **Figure S6A**  **Quantified** | Wild type | | | Day 5 |  |  |  |  | **21** | | | **0** | | | **21** | | **0.0** | |
|  |  |  |  | Day 10 |  |  |  |  | 70 | | | 3 | | | 73 | | 4.1 | |
|  |  |  |  | Day 15 |  |  |  |  | 19 | | | 20 | | | 39 | | 51.3 | |

**Supplementary Table 2: Details of *C. elegans* lifespan assays performed at 20ºC**

| **Lifespan assays performed post Day10-Smurf assay- Related to Figure 1E** | | | | | | | |
| --- | --- | --- | --- | --- | --- | --- | --- |
| **Strain identifier** | **Genotype** | **Food** | **Non-Smurfs/ Smurfs** | **Mean lifespan (MLS) days** | **% MLS w.r.t. non-Smurf** | ***P-value*** | **Animals Scored/Total** |
| Wild type  (Repeat 1) | N2-CGC | OP50 | **Non-Smurfs** | **19.2** |  |  | **17** |
|  |  |  | Smurfs | 15.8 | -17.7 | *0.0005* | 6 |
| Wild type  (Repeat 2)  ***Figure 1E*** | N2-CGC |  | **Non-Smurfs** | **20.1** |  |  | **13/14** |
|  |  |  | Smurfs | 15.7 | -21.8 | *<0.0001* | 18/19 |
| **Repeats for lifespan assays- Related to Figure 1, 2, and 3. RNAi was initiated on Day 1 of adulthood** | | | | | | | |
| **Strain identifier** | **Genotype** | **RNAi bacteria** | **Target tissue** | **Mean lifespan (MLS) days** | **% MLS w.r.t. control** | ***P-value*** | **Animals Scored/Total** |
| ***Lifespan data related to Figure 1F*** | | | | | | | |
| MAH677 (Repeat 1) ***Figure 1F*** | *sid-1(qt9) V; sqIs71[pMH1201/rgef-1p::gfp::unc-54 3’UTR+ pMH1141/rgef-1p::sid-1::unc-54 3'UTR + pMH477/pBS]* | control | Neuron-specific | **25.5** |  |  | **67/85** |
|  |  | *unc-31* |  | 29.1 | +14.1 | *<0.0001* | 75/89 |
| MAH346 (Repeat 1) | *sid-1(qt9) V* | control | RNAi-defective | **24.6** |  |  | **73/102** |
|  |  | *unc-31* |  | 24.5 | -0.5 | *0.6887* | 58/85 |
| MAH677 (Repeat 2) | *sid-1(qt9) V; sqIs71[pMH1201/rgef-1p::gfp::unc-54 3’UTR+ pMH1141/rgef-1p::sid-1::unc-54 3'UTR + pMH477/pBS]* | control | Neuron-specific | **20.3** |  |  | **85/124** |
|  |  | *unc-31* |  | 25.1 | +23.5 | *<0.0001* | 85/126 |
| MAH346 (Repeat 2) | *sid-1(qt9) V* | control | RNAi-defective | **22.8** |  |  | **75/119** |
|  |  | *unc-31* |  | 21.3 | -6.5 | *0.008* | 73/120 |
| MAH677 (Repeat 3) | *sid-1(qt9) V; sqIs71[pMH1201/rgef-1p::gfp::unc-54 3’UTR+ pMH1141/rgef-1p::sid-1::unc-54 3'UTR + pMH477/pBS]* | control | Neuron-specific | **22.8** |  |  | **87/122** |
|  |  | *unc-31* |  | 25.6 | +12.0 | *0.0015* | 85/120 |
| MAH346 (Repeat 3) | *sid-1(qt9) V* | control | RNAi-defective | **22.1** |  |  | **107/137** |
|  |  | *unc-31* |  | 22.1 | -0.1 | *0.9723* | 108/137 |
| ***Lifespan data related to Figure 1F*** | | | | | | | |
| MAH677 (Repeat 1) | *sid-1(qt9) V; sqIs71[pMH1201/rgef-1p::gfp::unc-54 3’UTR+ pMH1141/rgef-1p::sid-1::unc-54 3'UTR + pMH477/pBS]* | control | Neuron-specific | **22.2** |  |  | **74/115** |
|  |  | *egl-3* |  | 26.8 | +21.1 | *<0.0001* | 66/118 |
| MAH346 (Repeat 1) | *sid-1(qt9) V* | control | RNAi-defective | **24.4** |  |  | **68/98** |
|  |  | *egl-3* |  | 25.3 | +3.6 | *0.29* | 94/125 |
| MAH677 (Repeat 2) ***Figure 1F*** | *sid-1(qt9) V; sqIs71[pMH1201/rgef-1p::gfp::unc-54 3’UTR+ pMH1141/rgef-1p::sid-1::unc-54 3'UTR + pMH477/pBS]* | control | Neuron-specific | **25.5** |  |  | **67/85** |
|  |  | *egl-3* |  | 29.8 | +16.9 | *<0.0001* | 72/102 |
| MAH346 (Repeat 2) | *sid-1(qt9) V* | control | RNAi-defective | **24.6** |  |  | **72/103** |
|  |  | *egl-3* |  | 23.7 | -3.8 | *0.2051* | 54/82 |
| MAH677 (Repeat 3) | *sid-1(qt9) V; sqIs71[pMH1201/rgef-1p::gfp::unc-54 3’UTR+ pMH1141/rgef-1p::sid-1::unc-54 3'UTR + pMH477/pBS]* | control | Neuron-specific | **20.9** |  |  | **67/99** |
|  |  | *egl-3* |  | 23.6 | +12.9 | *0.0215* | 60/103 |
| MAH346 (Repeat 3) | *sid-1(qt9) V* | control | RNAi-defective | **21.3** |  |  | **49/115** |
|  |  | *egl-3* |  | 21.3 | 0.0 | *0.9005* | 54/112 |
| MAH677 (Repeat 4) | *sid-1(qt9) V; sqIs71[pMH1201/rgef-1p::gfp::unc-54 3’UTR+ pMH1141/rgef-1p::sid-1::unc-54 3'UTR + pMH477/pBS]* | control | Neuron-specific | **20.3** |  |  | **85/124** |
|  |  | *egl-3* |  | 24.4 | +20.1 | *0.0002* | 59/123 |
| MAH346 (Repeat 4) | *sid-1(qt9) V* | control | RNAi-defective | **22.8** |  |  | **75/119** |
|  |  | *egl-3* |  | 21.7 | -4.8 | *0.0788* | 71/129 |
| MAH677 (Repeat 5) | *sid-1(qt9) V; sqIs71[pMH1201/rgef-1p::gfp::unc-54 3’UTR+ pMH1141/rgef-1p::sid-1::unc-54 3'UTR + pMH477/pBS]* | control | Neuron-specific | **22.8** |  |  | **87/122** |
|  |  | *egl-3* |  | 28.1 | +23.2 | *<0.0001* | 82/128 |
| MAH346 (Repeat 5) | *sid-1(qt9) V* | control | RNAi-defective | **22.2** |  |  | **107/137** |
|  |  | *egl-3* |  | 22.4 | +0.9 | *0.9639* | 121/147 |
| ***Lifespan data related to Figure 2F*** | | | | | | | |
| MAH677 (Repeat 1)  ***Figure 2F*** | *sid-1(qt9) V; sqIs71[pMH1201/rgef-1p::gfp::unc-54 3’UTR+ pMH1141/rgef-1p::sid-1::unc-54 3'UTR + pMH477/pBS]* | control | Neuron-specific | **20.3** |  |  | **85/124** |
|  |  | *daf-28* |  | 23.4 | +15.1 | *0.002* | 73/123 |
| MAH346 (Repeat 1) | *sid-1(qt9) V* | control | RNAi-defective | **22.8** |  |  | **75/119** |
|  |  | *daf-28* |  | 21.9 | -4.3 | *0.1816* | 55/112 |
| MAH677 (Repeat 2) | *sid-1(qt9) V; sqIs71[pMH1201/rgef-1p::gfp::unc-54 3’UTR+ pMH1141/rgef-1p::sid-1::unc-54 3'UTR + pMH477/pBS]* | control | Neuron-specific | **22.8** |  |  | **87/122** |
|  |  | *daf-28* |  | 23.6 | +3.5 | *0.2294* | 101/135 |
| MAH346 (Repeat 2) | *sid-1(qt9) V* | control | RNAi-defective | **22.2** |  |  | **107/137** |
|  |  | *daf-28* |  | 22.7 | +2.6 | *0.6098* | 120/139 |
| ***Lifespan data related to Figure 2G*** | | | | | | | |
| MAH677 (Repeat 1) ***Figure 2G*** | *sid-1(qt9) V; sqIs71[pMH1201/rgef-1p::gfp::unc-54 3’UTR+ pMH1141/rgef-1p::sid-1::unc-54 3'UTR + pMH477/pBS]* | control | Neuron-specific | **21.2** |  |  | **76/122** |
|  |  | *ins-7* |  | 27.7 | +31.0 | *<0.0001* | 75/119 |
| MAH346 (Repeat 1) | *sid-1(qt9) V* | control | RNAi-defective | **21.3** |  |  | **71/119** |
|  |  | *ins-7* |  | 22.1 | +3.9 | *0.3576* | 58/123 |
| MAH677 (Repeat 2) | *sid-1(qt9) V; sqIs71[pMH1201/rgef-1p::gfp::unc-54 3’UTR+ pMH1141/rgef-1p::sid-1::unc-54 3'UTR + pMH477/pBS]* | control | Neuron-specific | **20.3** |  |  | **85/124** |
|  |  | *ins-7* |  | 27.8 | +36.4 | *<0.0001* | 88/117 |
| MAH346 (Repeat 2) | *sid-1(qt9) V* | control | RNAi-defective | **22.8** |  |  | **75/119** |
|  |  | *ins-7* |  | 22.4 | -2.0 | *0.5234* | 66/119 |
| MAH677 (Repeat 3) | *sid-1(qt9) V; sqIs71[pMH1201/rgef-1p::gfp::unc-54 3’UTR+ pMH1141/rgef-1p::sid-1::unc-54 3'UTR + pMH477/pBS]* | control | Neuron-specific | **22.8** |  |  | **87/122** |
|  |  | *ins-7* |  | 27.6 | +20.8 | *<0.0001* | 105/139 |
| MAH346 (Repeat 3) | *sid-1(qt9) V* | control | RNAi-defective | 22.2 |  |  | 107/137 |
|  |  | *ins-7* |  | 24.2 | +9.1 | *0.0019* | 113/136 |
| ***Lifespan data related to Figure 3C*** | | | | | | | |
| MAH728 (Repeat 1) | *sid-1(qt9) V; alxIs6[vha-6p::sid-1::sl2::gfp]* | control | Intestine-specific | **18.1** |  |  | **34/107** |
|  |  | *daf-2* |  | 29.1 | +60.7 | *<0.0001* | 36/99 |
| MAH346 (Repeat 1) | *sid-1(qt9) V* | control | RNAi-defective | **21.6** |  |  | **83/101** |
|  |  | *daf-2* |  | 26.0 | +20.3 | *<0.0001* | 79/96 |
| MAH728 (Repeat 2)  ***Figure 3C*** | *sid-1(qt9) V; alxIs6[vha-6p::sid-1::sl2::gfp]* | control | Intestine-specific | **24.0** |  |  | **89/110** |
|  |  | *daf-2* |  | 39.4 | +63.9 | *<0.0001* | 67/95 |
| MAH346 (Repeat 2) | *sid-1(qt9) V* | control | RNAi-defective | **24.4** |  |  | **68/98** |
|  |  | *daf-2* |  | 26.4 | +7.9 | *0.0105* | 75/113 |
| MAH728 (Repeat 3) | *sid-1(qt9) V; alxIs6[vha-6p::sid-1::sl2::gfp]* | control | Intestine-specific | **21.7** |  |  | **67/102** |
|  |  | *daf-2* |  | 35.6 | +64.2 | *<0.0001* | 62/99 |
| MAH346 (Repeat 3) | *sid-1(qt9) V* | control | RNAi-defective | **20.5** |  |  | **66/103** |
|  |  | *daf-2* |  | 25.6 | +25.2 | *<0.0001* | 57/102 |
| MAH728 (Repeat 4) | *sid-1(qt9) V; alxIs6[vha-6p::sid-1::sl2::gfp]* | control | Intestine-specific | **20.9** |  |  | **53/107** |
|  |  | *daf-2* |  | 33.3 | +59.5 | *<0.0001* | 58/123 |
| MAH346 (Repeat 4) | *sid-1(qt9) V* | control | RNAi-defective | **21.3** |  |  | **71/119** |
|  |  | *daf-2* |  | 22.1 | +3.8 | *0.1622* | 78/126 |
| MAH728 (Repeat 5) | *sid-1(qt9) V; alxIs6[vha-6p::sid-1::sl2::gfp]* | control | Intestine-specific | **21.2** |  |  | **49/111** |
|  |  | *daf-2* |  | 31.7 | +50.1 | *<0.0001* | 61/109 |
| MAH346 (Repeat 5) | *sid-1(qt9) V* | control | RNAi-defective | **20.0** |  |  | **58/111** |
|  |  | *daf-2* |  | 22.1 | +11.0 | *0.0071* | 49/111 |
| MAH728 (Repeat 6) | *sid-1(qt9) V; alxIs6[vha-6p::sid-1::sl2::gfp]* | control | Intestine-specific | **22.5** |  |  | **111/140** |
|  |  | *daf-2* |  | 36.2 | +61.1 | *<0.0001* | 105/140 |
| MAH346 (Repeat 6) | *sid-1(qt9) V* | control | RNAi-defective | **22.2** |  |  | **107/137** |
|  |  | *daf-2* |  | 24.1 | +8.7 | *0.0003* | 120/137 |
| ***Lifespan data related to Figure S4B*** | | | | | | | |
| MAH728 (Repeat 1)  ***Figure S4B*** | *sid-1(qt9) V; alxIs6[vha-6p::sid-1::sl2::gfp]* | control | Intestine-specific | **24.0** |  |  | **89/110** |
|  |  | *age-1* |  | 33.6 | +39.7 | *<0.0001* | 89/114 |
| MAH346 (Repeat 1) | *sid-1(qt9) V* | control | RNAi-defective | **24.4** |  |  | **68/98** |
|  |  | *age-1* |  | 28.3 | +16.1 | *<0.0001* | 62/98 |
| MAH728 (Repeat 2) | *sid-1(qt9) V; alxIs6[vha-6p::sid-1::sl2::gfp]* | control | Intestine-specific | **22.4** |  |  | **71/102** |
|  |  | *age-1* |  | 29.1 | +30.1 | *<0.0001* | 51/93 |
| MAH346 (Repeat 2) | *sid-1(qt9) V* | control | RNAi-defective | **24.6** |  |  | **73/102** |
|  |  | *age-1* |  | 30.2 | +22.7 | *<0.0001* | 41/82 |
| MAH728 (Repeat 3) | *sid-1(qt9) V; alxIs6[vha-6p::sid-1::sl2::gfp]* | control | Intestine-specific | **21.7** |  |  | **67/102** |
|  |  | *age-1* |  | 30.9 | +42.4 | *<0.0001* | 55/101 |
| MAH346 (Repeat 3) | *sid-1(qt9) V* | control | RNAi-defective | **20.5** |  |  | **66/103** |
|  |  | *age-1* |  | 26.0 | +27.1 | *<0.0001* | 63/104 |

**Supplementary Table 3: Details of Female *Drosophila* lifespans and Smurf assays**

| **Repeats for lifespan assays- Related to Figure 1, 2 and 3. Tissue-specific GAL-4 was**  **activated in 1–2-day old flies for 10 days at 29ºC. Flies were scored for lifespan day 5 onwards.** | | | | | | | | | |
| --- | --- | --- | --- | --- | --- | --- | --- | --- | --- |
| **Genotype** | **Target tissue** | **MLS (days)** | **% MLS w.r.t. Gal4/+** | ***P-value*** | **% MLS w.r.t.**  **UAS-RNAi/+** | ***P-value*** | **Total**  **flies** | **Cumulative Smurf%** | |
| ***Data related to Figure 1H (only lifespan)*** | | | | | | | | | |
| *elavC155-Gal4; TubGal80 / +*  (Repeat 1) | Neuronal GAL4  control | 45.8 |  |  |  |  | 57 | N.D. | |
| *elavC155-Gal4; TubGal80/UAS-RNAi-****cadps***  (Repeat 1) | Neuron-specific RNAi | **51.9** | **+13.3** | ***0.0002*** | **+21.3** | ***<0.0001*** | **120** |  |  |
| *UAS-RNAi-****cadps****/+*  (Repeat 1) | RNAi  control | 42.7 |  |  |  |  | 90 |  |  |
| *elavC155-Gal4; TubGal80 / +*  (Repeat 2) | Neuronal GAL4  control | 52.0 |  |  |  |  | 88 | N.D. | |
| *elavC155-Gal4; TubGal80/UAS-RNAi-****cadps***  (Repeat 2) | Neuron-specific RNAi | **56.3** | **+8.5** | ***0.0035*** | **+5.1** | ***0.2857*** | **40** |  |  |
| *UAS-RNAi-****cadps****/+*  (Repeat 2) | RNAi  control | 53.6 |  |  |  |  | 30 |  |  |
| ***Data related to Figure 1H and 1I (Lifespan + Smurf)*** | | | | | | | | **~Day 40 (Mid-life)** | **~Day 55 (old age)** |
| *elavC155-Gal4; TubGal80 / +*  (Repeat 3) | Neuronal GAL4  control | 42.0 |  |  |  |  | 83 | 10.8 | 10.8 |
| *elavC155-Gal4; TubGal80/UAS-RNAi-****cadps***  (Repeat 3) | Neuron-specific RNAi | **45.9** | **+9.1** | ***0.0195*** | **+28.8** | ***<0.0001*** | **78** | 1.3 | 3.9 |
| *UAS-RNAi-****cadps****/+*  (Repeat 3) | RNAi  control | 35.6 |  |  |  |  | 59 | 8.5 | 10.2 |
| *elavC155-Gal4; TubGal80 / +*  (Repeat 4)  ***Figure 1H*** | Neuronal GAL4  control | 53.3 |  |  |  |  | 60 | 5.1 | 18.7 |
| *elavC155-Gal4; TubGal80/UAS-RNAi-****cadps***  (Repeat 4) ***Figure 1H*** | Neuron-specific RNAi | **58.5** | **+9.5** | ***0.0033*** | **+17.8** | ***<0.0001*** | 48 | 2.1 | 8.5 |
| *UAS-RNAi-****cadps****/+*  (Repeat 4) | RNAi  control | 49.6 |  |  |  |  | 34 | 0 | 20.5 |
| ***Data related to Figure 1H (only lifespan)*** | | | | | | | | | |
| *elavC155-Gal4; TubGal80 / +*  (Repeat 1) | Neuronal GAL4  control | 45.8 |  |  |  |  | 57 | N.D. | |
| *elavC155-Gal4; TubGal80/UAS-RNAi-****amon***  (Repeat 1) | Neuron-specific RNAi | **51.2** | **+11.9** | ***0.0004*** | **+17.0** | ***0.0004*** | **105** |  |  |
| *UAS-RNAi-****amon****/+*  (Repeat 1) | RNAi  control | 43.8 |  |  |  |  | 74 |  |  |
| *elavC155-Gal4; TubGal80 / +*  (Repeat 2) | Neuronal GAL4  control | 52.0 |  |  |  |  | 88 | N.D. | |
| *elavC155-Gal4; TubGal80/UAS-RNAi-****amon***  (Repeat 2) | Neuron-specific RNAi | **55.2** | **+6.3** | ***0.0009*** | **-1.9** | ***0.7913*** | **80** |  |  |
| *UAS-RNAi-****amon****/+*  (Repeat 2) | RNAi  control | 56.3 |  |  |  |  | 23 |  |  |
| ***Data related to Figure 1H and 1J (Lifespan + Smurf)*** | | | | | | | | **~Day 40 (Mid-life)** | **~Day 55 (old age)** |
| *elavC155-Gal4; TubGal80 / +*  (Repeat 3) | Neuronal GAL4  control | 42.0 |  |  |  |  | 83 | 10.8 | 10.8 |
| *elavC155-Gal4; TubGal80/UAS-RNAi-****amon***  (Repeat 3) | Neuron-specific RNAi | **43.2** | **+2.8** | ***0.1646*** | **-0.8** | ***0.4017*** | **51** | **0** | **0** |
| *UAS-RNAi-****amon*** */+*  (Repeat 3) | RNAi  control | 43.5 |  |  |  |  | 114 | 9.7 | 13.2 |
| *elavC155-Gal4; TubGal80 / +*  (Repeat 4) | Neuronal GAL4  control | 45.7 |  |  |  |  | 50 | 2.6 | 15.7 |
| *elavC155-Gal4; TubGal80/UAS-RNAi-****amon***  (Repeat 4) | Neuron-specific RNAi | **51.0** | **+11.7** | ***0.2893*** | **+30.2** | ***0.2472*** | **26** | **0** | **0** |
| *UAS-RNAi-****amon****/+*  (Repeat 4) | RNAi  control | 39.2 |  |  |  |  | 58 | 4.7 | 11.9 |
| *elavC155-Gal4; TubGal80 / +*  (Repeat 5)  ***Figure 1H*** | Neuronal GAL4  control | 53.3 |  |  |  |  | 60 | 5.1 | 18.7 |
| *elavC155-Gal4; TubGal80/UAS-RNAi-****amon***  (Repeat 5) ***Figure 1H*** | Neuron-specific RNAi | **58.5** | **+9.7** | ***0.0072*** | **+5.3** | ***0.166*** | **60** | **5** | **13.3** |
| *UAS-RNAi-****amon****/+*  (Repeat 5) | RNAi  control | 55.6 |  |  |  |  | 40 | 1.6 | 31.6 |
| ***Data related to Figure 2 (Lifespan + Smurf)*** | | | | | | | | **~Day 40 (Mid life)** | **~Day 55 (old age)** |
| *elavC155-Gal4; TubGal80/UAS-****GFP***  (Repeat 1) | Neuronal GFP  control | 32.2 |  |  |  |  | 60 | 15.8 | 26.3 |
| *elavC155-Gal4; TubGal80/UAS-RNAi-****dilp2***  (Repeat 1) | Neuron-specific RNAi | 39.6 | +22.8 |  | N.D. | N.D. | 30 | 0 | 0 |
| *elavC155-Gal4; TubGal80/UAS-RNAi-****dilp3***  (Repeat 1) | Neuron-specific RNAi | 42.4 | +31.6 | *0.0014* | N.D. | N.D. | 94 | 0 | 0 |
| *elavC155-Gal4; TubGal80/UAS-RNAi-****dilp5***  (Repeat 1) | Neuron-specific RNAi | 39.4 | +22.5 | *0.0087* | N.D. | N.D. | 79 | 0 | 2.6 |
| ***Data related to Figure 2H (only lifespan)*** | | | | | | | | | |
| *elavC155-Gal4; TubGal80 / +*  (Repeat 1) | Neuronal GAL4  control | 52.0 |  |  |  |  | 88 | N.D. | |
| *elavC155-Gal4; TubGal80/UAS-RNAi-****dilp3***  (Repeat 1) | Neuron-specific RNAi | **58.5** | **+12.7** | ***<0.0001*** | **+9.4** | ***0.0133*** | **79** |  |  |
| *UAS-RNAi-****dilp3****/+*  (Repeat 1) | RNAi  control | 53.5 |  |  |  |  | 62 |  |  |
| ***Data related to Figure 2H (Lifespan + Smurf)*** | | | | | | | | **~Day 40 (Mid-life)** | **~Day 55 (old age)** |
| *elavC155-Gal4; TubGal80 / +*  (Repeat 2)  ***Figure 2H*** | Neuronal GAL4  control | 53.3 |  |  |  |  | 60 | 5.1 | 18.7 |
| *elavC155-Gal4; TubGal80/UAS-RNAi-****dilp3***  (Repeat 2)  ***Figure 2H*** | Neuron-specific RNAi | **59.2** | **+11.1** | ***0.001*** | **+14.1** | ***0.0267*** | 97 | 4.2 | 10.6 |
| *UAS-RNAi-****dilp3****/+*  (Repeat 2) | RNAi  control | 51.9 |  |  |  |  | 20 | 0 | 15 |
| ***Data related to Figure 3D (only lifespan)*** | | | | | | | | | |
| *Np1-Gal4; TubGal80 / +* (Repeat 1) | Intestinal GAL4  control | 52.0 |  |  |  |  | 45 | N.D. | |
| *Np1-Gal4; TubGal80/*  *UAS-RNAi-****InR***  (Repeat 1) | Intestine-specific RNAi | **62.1** | **+19.5** | ***<0.0001*** | **+10.0** | ***0.0029*** | **37** |  |  |
| *UAS-RNAi-****InR****/+*  (Repeat 1) | RNAi  control | 57.3 |  |  |  |  | 31 |  |  |
| *Np1-Gal4; TubGal80 / +* (Repeat 2) | Intestinal GAL4  control | 52.3 |  |  |  |  | 45 | N.D. | |
| *Np1-Gal4; TubGal80/*  *UAS-RNAi-****InR***  (Repeat 2) | Intestine-specific RNAi | **58.0** | **+10.8** | ***<0.0001*** | **+7.9** | ***0.0006*** | **45** |  |  |
| *UAS-RNAi-****InR****/+*  (Repeat 2) | RNAi  control | 53.7 |  |  |  |  | 45 |  |  |
| ***Data related to Figure 3D (Lifespan + Smurf)*** | | | | | | | | **~Day 40 (Mid-life)** | **~Day 55 (old age)** |
| *Np1-Gal4; TubGal80 / +* (Repeat 4)  ***Figure 3D*** | Intestinal GAL4  control | 51.0 |  |  |  |  | 58 | N.D. | 17.2 |
| *Np1-Gal4; TubGal80/*  *UAS-RNAi-****InR***  (Repeat 4) ***Figure 3D*** | Intestine-specific RNAi | **55.0** | **+8.0** | ***0.0003*** | **+4.6** | ***0.0031*** | **54** | **N.D.** | **13.0** |
| *UAS-RNAi-****InR****/+*  (Repeat 4) | RNAi  control | 52.6 |  |  |  |  | 61 | N.D. | 18.0 |

**Supplementary Table 4: *C. elegans* Smurf screen for neuron-specific RNAi of insulin-like peptides (ILPs)**

| **Insulin Class** | **Strain identifier** | **Target tissue** | **RNAi bacteria** | **# of Non-Smurf** | **# of Smurf** | **Total # of animals** | **% Smurf** | **Change in % Smurf** |
| --- | --- | --- | --- | --- | --- | --- | --- | --- |
| **Type-Gamma** | **MAH677** | Neuron-specific  RNAi | **control** | **17** | **42** | **59** | **71.2** |  |
|  |  |  | *ins-11* | 22 | 31 | 53 | 58.5 | -12.7 |
|  |  |  | **control** | **4** | **24** | **28** | **85.7** |  |
|  |  |  | *ins-11* | 4 | 15 | 19 | 78.9 | - 6.8 |
|  |  |  | **control** | **19** | **10** | **29** | **34.5** |  |
|  |  |  | *ins-12* | 25 | 26 | 51 | 51.0 | +16.5 |
|  |  |  | *ins-14* | 15 | 17 | 32 | 53.0 | +18.6 |
|  |  |  | *ins-15* | 14 | 14 | 28 | 50.0 | +15.5 |
|  |  |  | **control** | **12** | **13** | **25** | **52.0** |  |
|  |  |  | *ins-16* | 10 | 4 | 14 | 28.6 | -23.4 |
|  |  |  | **control** | **9** | **12** | **21** | **57.1** |  |
|  |  |  | *ins-16* | 10 | 6 | 16 | 37.5 | -19.6 |
|  |  |  | **control** | **3** | **26** | **29** | **89.7** |  |
|  |  |  | *ins-17* | 4 | 23 | 27 | 85.2 | - 4.5 |
|  |  |  | *ins-18* | 3 | 22 | 25 | 88.0 | - 1.7 |
|  |  |  | **control** | **4** | **27** | **31** | **87.1** |  |
|  |  |  | *ins-17* | 3 | 43 | 46 | 93.5 | + 6.4 |
|  |  |  | *ins-18* | 5 | 23 | 28 | 82.1 | - 5.0 |
|  |  |  | **control** | **3** | **30** | **33** | **90.9** |  |
|  |  |  | *ins-32* | 8 | 25 | 33 | 75.8 | -15.2 |
|  |  |  | *ins-37* | 14 | 26 | 40 | 65.0 | -25.9 |
|  |  |  | **control** | **4** | **16** | **20** | **80.0** |  |
|  |  |  | *ins-32* | 5 | 21 | 26 | 80.8 | + 0.8 |
|  |  |  | *ins-37* | 6 | 32 | 38 | 84.2 | + 4.2 |
|  |  |  | **control** | **3** | **12** | **15** | **80.0** |  |
|  |  |  | *ins-37* | 5 | 10 | 15 | 66.7 | -13.3 |
| **Type-**  **Beta** | **MAH677** | Neuron-specific  RNAi | **control** | **17** | **42** | **59** | **71.2** |  |
|  |  |  | *ins-1* | 15 | 24 | 39 | 61.5 | - 9.6 |
|  |  |  | *ins-2* | 27 | 40 | 67 | 59.7 | -11.5 |
|  |  |  | *ins-3* | 26 | 34 | 60 | 56.7 | -14.5 |
|  |  |  | *ins-4* | 21 | 35 | 56 | 62.5 | - 8.7 |
|  |  |  | *ins-5* | 32 | 35 | 67 | 52.2 | - 18.9 |
|  |  |  | *ins-6* | 20 | 27 | 47 | 57.4 | -13.7 |
|  |  |  | *ins-7* | 39 | 19 | 58 | 32.8 | -38.4 |
|  |  |  | **control** | **4** | **24** | **28** | **85.7** |  |
|  |  |  | *ins-1* | 2 | 19 | 21 | 90.5 | + 4.8 |
|  |  |  | *ins-2* | 4 | 24 | 28 | 85.7 | 0.0 |
|  |  |  | *ins-3* | 6 | 13 | 19 | 68.4 | -17.3 |
|  |  |  | *ins-4* | 1 | 16 | 17 | 94.1 | + 8.4 |
|  |  |  | *ins-5* | 6 | 28 | 34 | 82.4 | - 3.4 |
|  |  |  | *ins-6* | 5 | 15 | 20 | 75.0 | -10.7 |
|  |  |  | *ins-7* | 9 | 22 | 31 | 71.0 | -14.7 |
|  |  |  | **control** | **8** | **8** | **16** | **50.0** |  |
|  |  |  | *ins-1* | 12 | 11 | 23 | 47.8 | - 2.2 |
|  |  |  | *ins-2* | 8 | 14 | 22 | 63.6 | +13.6 |
|  |  |  | *ins-3* | 11 | 6 | 17 | 35.3 | -14.7 |
|  |  |  | *ins-4* | 8 | 13 | 21 | 61.9 | +11.9 |
|  |  |  | *ins-5* | 10 | 10 | 20 | 50.0 | 0.0 |
|  |  |  | *ins-6* | 22 | 12 | 34 | 35.3 | -14.7 |
|  |  |  | **control** | **13** | **25** | **38** | **65.8** |  |
|  |  |  | *ins-5* | 6 | 27 | 33 | 81.8 | +16.0 |
|  |  |  | *ins-7* | 18 | 11 | 29 | 37.9 | -27.9 |
|  |  |  | **control** | **20** | **56** | **76** | **73.7** |  |
|  |  |  | *ins-7* | 39 | 19 | 58 | 32.8 | -40.9 |
|  |  |  | **control** | **19** | **10** | **29** | **34.5** |  |
|  |  |  | *ins-9* | 18 | 15 | 33 | 45.5 | +11.0 |
|  |  |  | **control** | **20** | **56** | **76** | **73.7** |  |
|  |  |  | *daf-28* | 31 | 29 | 60 | 48.3 | -25.4 |
|  |  |  | **control** | **17** | **34** | **51** | **66.7** |  |
|  |  |  | *daf-28* | 39 | 34 | 73 | 46.6 | -20.1 |
| **Type-**  **Alpha** | **MAH677** | Neuron-specific  RNAi | **control** | **12** | **13** | **25** | **52.0** |  |
|  |  |  | *ins-20* | 17 | 12 | 29 | 41.4 | -10.6 |
|  |  |  | **control** | **3** | **26** | **29** | **89.7** |  |
|  |  |  | *ins-21* | 4 | 11 | 15 | 73.3 | -16.3 |
|  |  |  | *ins-22* | 4 | 11 | 15 | 73.3 | -16.3 |
|  |  |  | *ins-23* | 9 | 13 | 22 | 59.1 | -30.6 |
|  |  |  | **control** | **4** | **27** | **31** | **87.1** |  |
|  |  |  | *ins-21* | 6 | 39 | 45 | 86.7 | - 0.4 |
|  |  |  | *ins-22* | 14 | 32 | 46 | 69.6 | -17.5 |
|  |  |  | *ins-23* | 9 | 14 | 23 | 60.9 | -26.2 |
|  |  |  | **control** | **4** | **12** | **16** | **75.0** |  |
|  |  |  | *ins-21* | 8 | 34 | 42 | 81.0 | + 6.0 |
|  |  |  | *ins-22* | 9 | 27 | 36 | 75.0 | 0.0 |
|  |  |  | *ins-23* | 3 | 19 | 22 | 86.4 | +11.4 |
|  |  |  | **control** | **12** | **13** | **25** | **52.0** |  |
|  |  |  | *ins-24* | 12 | 14 | 26 | 53.8 | + 1.8 |
|  |  |  | *ins-25* | 22 | 15 | 37 | 40.5 | -11.5 |
|  |  |  | **control** | **3** | **26** | **29** | **89.7** |  |
|  |  |  | *ins-26* | 6 | 16 | 22 | 72.7 | -16.9 |
|  |  |  | **control** | **4** | **27** | **31** | **87.1** |  |
|  |  |  | *ins-26* | 8 | 25 | 33 | 75.8 | -11.3 |
|  |  |  | **control** | **12** | **13** | **25** | **52.0** |  |
|  |  |  | *ins-27* | 18 | 8 | 26 | 30.8 | -21.2 |
|  |  |  | **control** | **4** | **53** | **57** | **93.0** |  |
|  |  |  | *ins-27* | 14 | 40 | 54 | 74.1 | -18.9 |
|  |  |  | **control** | **9** | **12** | **21** | **57.1** |  |
|  |  |  | *ins-28* | 16 | 4 | 20 | 20.0 | -37.1 |
|  |  |  | *ins-29* | 10 | 6 | 16 | 37.5 | -19.6 |
|  |  |  | **control** | **3** | **26** | **29** | **89.7** |  |
|  |  |  | *ins-30* | 10 | 20 | 30 | 66.7 | -23.0 |
|  |  |  | **control** | **4** | **27** | **31** | **87.1** |  |
|  |  |  | *ins-30* | 4 | 25 | 29 | 86.2 | - 0.9 |
|  |  |  | **control** | **3** | **12** | **15** | **80.0** |  |
|  |  |  | *ins-33* | 7 | 15 | 22 | 68.2 | -11.8 |
|  |  |  | *ins-35* | 5 | 9 | 14 | 64.3 | -15.7 |
|  |  |  | **control** | **3** | **30** | **33** | **90.9** |  |
|  |  |  | *ins-33* | 5 | 24 | 29 | 82.8 | - 8.2 |
|  |  |  | *ins-35* | 6 | 18 | 24 | 75.0 | -15.9 |
|  |  |  | **control** | **4** | **16** | **20** | **80.0** |  |
|  |  |  | *ins-33* | 12 | 14 | 26 | 53.8 | -26.2 |
|  |  |  | *ins-35* | 4 | 27 | 31 | 87.1 | + 7.1 |
|  |  |  | **control** | **9** | **12** | **21** | **57.1** |  |
|  |  |  | *ins-38* | 14 | 12 | 26 | 46.2 | -11.0 |
|  |  |  | **control** | **3** | **12** | **15** | **80.0** |  |
|  |  |  | *ins-39* | 7 | 17 | 24 | 70.8 | - 9.2 |
|  |  |  | **control** | **4** | **16** | **20** | **80.0** |  |
|  |  |  | *ins-39* | 8 | 22 | 30 | 73.3 | - 6.7 |
|  |  |  | **control** | **3** | **30** | **33** | **90.9** |  |
|  |  |  | *ins-39* | 13 | 28 | 41 | 68.3 | -22.6 |
| **Multiple chains** | **MAH677** | Neuron-specific  RNAi | **control** | **3** | **26** | **29** | **89.7** |  |
|  |  |  | *ins-31* | 8 | 14 | 22 | 63.6 | -26.0 |
|  |  |  | **control** | **4** | **27** | **31** | **87.1** |  |
|  |  |  | *ins-31* | 6 | 31 | 37 | 83.8 | - 3.3 |
|  |  |  | **control** | **4** | **12** | **16** | **75.0** |  |
|  |  |  | *ins-31* | 5 | 15 | 20 | 75.0 | 0.0 |

**Supplementary Table 5: List of *C. elegans* strains used in this study**

| **Strain identifier** | **Genotype** | **Source** | **Description** |
| --- | --- | --- | --- |
| Wild type (WT) | Bristol N2 | Caenorhabditis Genetics Center (CGC) | Wild type |
| Wild type (WT) | Bristol N2 | Kenyon lab | Wild type |
| CF2222 | *muEx336[mtl-1::RFP + rol-6]* | CGC | *mtl-1::RFP* reporter |
| CF1934 | *daf-16(mu86)I; muIs109[daf-16p::gfp::daf-16 + odr-1p::rfp]* | Dr. Cynthia Kenyon (UCSF) | DAF-16::GFP fusion reporter |
| CF1903 | *glp-1(e2144ts)* | Dr. Cynthia Kenyon (UCSF) | temperature sensitive germline-less mutant |
| MAH677 | *sid-1(qt9) V; sqIs71[pMH1201/rgef-1p::gfp::unc-54 3’UTR+ pMH1141/rgef-1p::sid-1::unc-54 3'UTR + pMH477/pBS]* | Y. Yang, C. Kumsta and M. Hansen, manuscript in preparation | Neuron-specific RNAi transgenic |
| MAH346 | *sid-1(qt9)* | parent strain AGD09: *sid-1(qt9) V* was provided by Dr. Andrew Dillin (UC Berkeley) and 4X outcrossed to N2-CK | RNAi-defective mutant |
| MAH728 | *sid-1(qt9) V; alxIs6[vha-6p::sid-1::sl2::gfp]* | Parent strain  parent strain MGH167: *sid-1(qt9); alxIs6[vha-6p::sid-1::sl2::gfp]* was provided by Dr. Alexander Soukas (Massachusetts General Hospital) and 4X outcrossed to MAH346 | Intestine-specific RNAi transgenic |
| MAH867 | *daf-16(mu86)+/- I?; sid-1(qt9) V; sqEx131[pMH1141/rgef-1p::sid-1::unc-54 3’UTR + pMH876/unc-122p::rfp]; muIs109[daf-16p::gfp::daf-16 + odr-1p::rfp]* | was obtained by standard genetic crosses | DAF-16::GFP fusion reporter in neuron-specific RNAi background |
| MAH1172 | *sid-1(qt9) V; sqIs71[pMH1201/rgef-1p::gfp::unc-54 3’UTR+ pMH1141/rgef-1p::sid-1::unc-54 3’UTR + pMH477/pBS]; muEx336[mtl-1::RFP + rol-6]* | was obtained by standard genetic crosses | *mtl-1::RFP* reporter in neuron-specific RNAi background |
| MAH1173 | *sid-1(qt9) V; alxIs6[vha-6p::sid-1::sl2::gfp]; muEx336[mtl-1::RFP + rol-6]* | was obtained by standard genetic crosses | *mtl-1::RFP* reporter in intestine-specific RNAi background |
| MAH1094 | *sqEx206[pMH1342/rgef-1p::egl-3c cDNA::pSM::mCherry]* | generated in this study by injecting N2-CGC strain with 100ng/μl of  *rgef-1p::egl-3c::pSM::mCherry* | Neuronal over-expressor of *egl-3* |
| MAH1256 | *sqEx236[rgef-1p::pSM::mCherry]* | generated in this study by injecting N2-CGC strain with 80ng/μl *rgef-1p::pSM::mCherry* | Four independent lines of control strains for neuronal over-expressor of *daf-28, ins-7* etc. |
| MAH1257 | *sqEx237[rgef-1p::pSM::mCherry]* |  |  |
| MAH1258 | *sqEx238[rgef-1p::pSM::mCherry]* |  |  |
| MAH1259 | *sqEx239[rgef-1p::pSM::mCherry]* |  |  |
| MAH1264 | *sqEx243[rgef-1p::daf-28(SM)::mCherry]* | generated in this study by injecting N2-CGC strain with 80ng/μl of *rgef-1p::daf-28(SM)::mCherry* | two independent lines of neuronal over-expressors of *daf-28* |
| MAH1265 | *sqEx244[rgef-1p::daf-28(SM)::mCherry]* |  |  |
| MAH1260 | *sqEx240[rgef-1p::ins-7(2b) gDNA::mCherry]* | generated in this study by injecting N2-CGC strain with 80ng/μl of *rgef-1p::ins-7(2b) gDNA::mCherry* | three independent lines of neuronal over-expressors of *ins-7* |
| MAH1261 | *sqEx241[rgef-1p::ins-7(2b) gDNA::mCherry]* |  |  |
| MAH1262 | *sqEx242[rgef-1p::ins-7(2b) gDNA::mCherry]* |  |  |
| MAH1093 | *sid-1(qt9) V; ueEx415[h20p::unc-31 sense::sl2::mCherry + h20p::unc-31 antisense::sl2::mCherry + elt-2p::gfp]; alxIs6[vha-6p::sid-1::sl2::gfp]* | generated in this study by standard genetic crosses | Neuronal *unc-31* hairpin transgenics combined with intestinal feeding RNAi transgenics |
| KUM11 | *glp-1(e2144ts); sqEx238[rgef-1p::pSM::mCherry]* | generated in this study by standard genetic crosses | control strain for KUM12 |
| KUM12 | *glp-1(e2144ts); sqEx244[rgef-1p::daf-28(SM)::mCherry* | generated in this study by standard genetic crosses | neuronal over-expressor of *daf-28* in *glp-1(e2144ts)* background |
| MAH1227 | *svIs69 [daf-28p::daf-28::GFP + unc-4(+)]* | parent strain VB1605: *svIs69 [daf-28p::daf-28::GFP + unc-4(+)]* was obtained from CGC, 4X outcrossed to N2-CK. | DAF-28::GFP fusion reporter |

**Supplementary Table 6: List of RNAi bacteria used in this study**

| **RNAi bacteria** | **Source** |
| --- | --- |
| L4440 (empty vector) control | Dr. Andrew Fire |
| *daf-2* | Dr. Andrew Dillin |
| *age-1* | Dr. Andrew Dillin |
| *daf-16* | Dr. Andrew Dillin |
| *daf-28* | Dr. Arnab Mukhopadhyay (from Vidal library) |
| *unc-31* | This study |
| *unc-13* | Ahringer RNAi library |
| *snb-1* | Ahringer RNAi library |
| *kpc-1* | Ahringer RNAi library |
| *egl-3* | Ahringer RNAi library |
| *sbt-1* | Ahringer RNAi library |
| *ins-2* | Ahringer RNAi library |
| *ins-4* | Ahringer RNAi library |
| *ins-9* | Ahringer RNAi library |
| *ins-12* | Ahringer RNAi library |
| *ins-14* | Ahringer RNAi library |
| *ins-15* | Ahringer RNAi library |
| *ins-16* | Ahringer RNAi library |
| *ins-17* | Ahringer RNAi library |
| *ins-18* | Ahringer RNAi library |
| *ins-20* | Ahringer RNAi library |
| *ins-21* | Ahringer RNAi library |
| *ins-23* | Ahringer RNAi library |
| *ins-24* | Ahringer RNAi library |
| *ins-25* | Ahringer RNAi library |
| *ins-27* | Ahringer RNAi library |
| *ins-29* | Ahringer RNAi library |
| *ins-30* | Ahringer RNAi library |
| *ins-31* | Ahringer RNAi library |
| *ins-32* | Ahringer RNAi library |
| *ins-35* | Ahringer RNAi library |
| *ins-38* | Ahringer RNAi library |
| *ins-39* | Ahringer RNAi library |
| *act-5* | Ahringer RNAi library |
| *clc-1* | Ahringer RNAi library |
| *let-413* | Ahringer RNAi library |
| *ajm-1* | Ahringer RNAi library |
| *aex-5* | Vidal RNAi library |
| *bli-4* | Vidal RNAi library |
| *egl-21* | Vidal RNAi library |
| *ins-1* | Vidal RNAi library |
| *ins-3* | Vidal RNAi library |
| *ins-5* | Vidal RNAi library |
| *ins-6* | Vidal RNAi library |
| *ins-7* | Vidal RNAi library |
| *ins-11* | Vidal RNAi library |
| *ins-22* | Vidal RNAi library |
| *ins-26* | Vidal RNAi library |
| *ins-33* | Vidal RNAi library |
| *ins-37* | Vidal RNAi library |
